# Lipid Specificity of the Fusion of Bacterial Extracellular Vesicles with the Host Membrane

**DOI:** 10.1101/827519

**Authors:** Ashutosh Prince, Anuj Tiwari, Titas Mandal, Debraj Koiri, Geetanjali Meher, Deepak Kumar Sinha, Mohammed Saleem

## Abstract

Bacterial membrane vesicles (MVs) facilitate long-distance delivery of virulence factors crucial for pathogenicity. The entry and trafficking mechanisms of virulence factors inside host cells are recently emerging, however, if bacterial MVs can fuse and modulate the physicochemical properties of the host lipid membrane and membrane lipid parameter for fusion remains unknown. Here we reconstitute the interaction of bacterial MV with host cell lipid membranes and quantitatively show that bacterial MV interaction increases the fluidity, dipole potential, and Compressibility of a biologically relevant multi-component host membrane upon fusion. The presence of cylindrical lipids such as phosphatidylcholine and a moderate acyl chain length of C16 helps the MV interaction. While significant binding of bacterial MVs to the raft-like lipid membranes with phase-separated regions of the membrane was observed, however, MVs prefer binding to the liquid-disordered regions of the membrane. Further, the elevated levels of cholesterol tend to hinder the interaction of bacterial MVs as evident from the favorable excess Gibbs free energy of mixing of bacterial MVs with host lipid membranes. The findings provide new insights that might have implications for the regulation of host machinery by bacterial pathogens through the manipulation of host membrane properties.

## Introduction

The secretion of membrane vesicles (MVs) from the bacterial cells is recognized as a generalized phenomenon observed in many bacteria ^1^. Bacterial MVs are known to facilitate contact-free inter-species communication, nutrient acquisition, enhanced survival against immune response and host-pathogen interaction ^2^. Initially known to be only secreted by controlled blebbing of Gram negative bacteria, emerging evidence supports the existence of different types of Bacterial MV, such as – Bacterial extracellular vesicles (MVs), Outer-inner membrane vesicles (OIBMVs) and Explosive outer membrane vesicles (EOBMVs) all of which are involved in transport of virulence factors ^3^. Typically, bacterial MVs are nano-sized spherical membrane compartments spanning 20-300 nm in diameter derived from the outer membrane lipid bilayer and the periplasm of the bacteria ^2^. Biochemically, bacterial MVs are known to carry lipopolysaccharides (LPS), phospholipids, peptidoglycans, outer membrane proteins (OMPs), nucleic acids, signaling molecules and periplasmic, cytoplasmic and membrane-bound proteins ^4^. The degree of bacterial MV secretion has also been correlated to the pathogenicity and virulence of bacteria ^5^. Interestingly, in addition to delivering a selection of virulence factors and immunomodulatory molecules directly into host cells during infection, bacterial MVs are also known to act as decoy antigen and divert the host immune system away from the bacterial cell ^6^. There is a considerable focus on the understanding of the virulence factors of the MV cargo and modes of bacterial MV entry inside host cells. Owing to a wide range of size of bacterial MVs, different pathways such as macropinocytosis, clathrin-dependent and caveolin or lipid raft mediated endocytosis have been implicated in the internalization processes of different Bacterial MVs ^7^. Further, direct membrane fusion with host cell membranes, preferentially at lipid raft domains, has also been reported to be a potential mechanism of entry of bacterial MVs ^8^. Similarly, fusion of bacterial MVs extracted from *L. pneumophila* with host model membranes suggested the faster time scales of fusion events (few seconds), which is also temperature-dependent ^9^.

A crucial aspect that remains unexplored is if/how MV fusion modulates the physicochemical parameters of host cell membranes (i.e., fluidity, dipole potential and in-plane Compressibility). Irrespective of the kind of bacterial MVs, mode of entry of bacterial MVs, insights on membrane interfacial interaction of bacterial MVs is crucial as the local lipid reorganization and remodeling of the host cell membrane is a prerequisite for MV entry. It is proposed that the MV fusion may result in changes in the properties of the host cell membrane ^9^, however, there is no experimental evidence in support of this idea. The lipid specificity, quantitative changes in physicochemical parameters (e.g., fluidity, dipole potential and in-plane Compressibility) of host membrane triggered by bacterial MVs and the thermodynamics of their interaction has never been explored to best of our knowledge.

Using *in vitro* reconstitution, confocal laser scanning microscopy, potentiometric dye-based fluorescence anisotropy, dipole potential measurements and surface pressure-area isotherms, we show that MV interaction increases the fluidity, dipole potential and in-plane Compressibility of the lipid membranes mimicking the host cell membrane. We also report the role of host membrane lipid head-group specificity, acyl chain length and phase separated boundaries on the interfacial interaction of bacterial MVs. The presence of lipids such as 1,2-dioleoyl-*sn*-glycero-3-phosphocholine (DOPC) and an acyl chain length of C16 are significant for the fusion of the bacterial MVs. Furthermore, the bacterial MVs prefer liquid disordered regions of the lipid membrane, whereas the elevated levels of cholesterol in raft-like lipid membranes with phase separated regions tend to hinder the interaction. The findings are further supported by the investigation of the thermodynamic parameters of MV interaction with host lipid membrane, revealing the change in Gibbs excess free energy of mixing of MV components with biologically relevant host membrane lipid membrane conditions. These findings show that bacterial membrane vesicles can effectively modulate the physicochemical properties of the host membrane that might eventually have consequences for membrane associated signaling during host-pathogen interactions.

## Materials and Methods

### Materials

Calcium chloride (CaCl_2_), HEPES, sodium chloride (NaCl), TX-100, Lauria Bertini (LB) broth, methanol, chloroform, Nile red were purchased from Himedia, India. Di-8-ANEPPS, Vybrant DiO were procured from ThermoFisher, USA. N-ethylmaleimide (NEM), Dimethylsulphoxide (DMSO) and RPMI media were purchased from Sigma Aldrich, USA. All lipids used in study, 1,2-dioleoyl-*sn*-glycero-3-phosphocholine (DOPC), 1,2-dilauroyl-sn-glycero-3-phosphocholine (DLPC), 1,2-dimyrisitol-sn-glycero-3-phophocholine (DMPC), 1,2-distearoyl-sn-glycero-3-phosphocholine (DSPC), 1,2-dipalmitoyl-sn-glycero-3-phosphocholine (DPPC), Sphingomyelin from Brain Porcine (BSM), cholesterol, 1,2-dioleoyl-sn-glycero-3-phosphoethanolamine (DOPE), 1,2-dioleoyl-sn-glycero-3-phophoethanolamine-N-(7-nitro-2-1,3-benzoxadiazol-4-yl) (NBD-PE), 1,2-dipalmitoyl-sn-glycero-3-phosphoethanolamine-N-(lissamine rhodamine B sulfonyl (Rhodamine-PE) were purchased from Avanti Polar lipids, USA. Monocytic leukemia cell lines (THP-1) were purchased from the National Center of Cell Science, Pune, India. *E. coli* used for isolation of extracellular vesicles was procured from Institute of Microbial Technology, Chandigarh, India.

### Preparation of Buffers and Solutions

Membrane vesicle (MV) buffer containing 10 mM HEPES, 2 mM CaCl_2_ and 150 mM NaCl prepared in deionized water with physiological pH of 7.4. 3mM N-ethylmaleimide (NEM) solution was made freshly during the experiment by dissolving NEM in miliQ water. Nile red powder was dissolved in DMSO (dimethylsulphoxide) to the final concentration of 100 nM. DiO labelling stock solution was prepared by dissolving 1mg DiO in 1 mL of DMSO.

### Membrane Vesicles Purification from Bacteria

*E. coli* were used for the vesiculation and isolation of bacterial membrane vesicles (MVs). The methodology for MV vesiculation and isolation was adopted from the procedure described by Sezgin et. al. ^10^ with some modification and optimization that allowed us to obtain reasonably homogenous pool of MVs having size in the range of 50-150 nm. We optimized the method that allowed us to isolate bacterial MVs without the usage of ultracentrifugation. However, we also isolated the bacterial MVs by using ultracentrifugation method to compare the MVs isolated by chemical vesiculation method. Briefly, bacterial cells were cultured in Luria Bertani (LB) broth by inoculating a single colony of *E. coli* from slant culture. The culture was incubated at 120 rpm at 37°C. Cells were grown until optical density (OD) of culture reached to a value of 1.0 where, considered as the *E. coli* are in mid-exponential phase of growth. Cells were harvested from media by centrifugation at 5000 rpm for 10 minutes at 4°C then supernatant were discarded. The pallet was re-suspended in 50 mL Membrane vesicle (MV) buffer (pH-7.4). Cells were again centrifuged at 5000 rpm for 10 minutes and washed twice in MV buffer to remove remaining media. Pellet of bacterial cells was then resuspended in 30 ml of 3mM of N-ethyl-maleimide (NEM) solution. N-ethyl maleimide (NEM) is well optimized chemical vesiculant for obtaining Giant Plasma Membrane Vesicles (GPMV) from eukaryotic cell membranes. The choice of NEM as a chemical vesiculant is justified by the fact that it is known not to cross-link with proteins and aldehydes or cause coupling of phosphatidylethanolamine to proteins or depalmitoylation. Resuspended cells were incubated in NEM containing MV buffer for 1 hour for vesiculation of bacterial MVs from *E. coli*. Vesiculated bacterial MVs in cells were isolated by high-speed centrifugation at 18000g for 2 hours at 4
°C and supernatant containing membrane vesicles (bacterial MVs) was collected. The bacterial MVs in supernatant was concentrated by using a 10 kD cut-off filter through 0.22 µm filter paper for the uniformity in size of bacterial MVs. For conventional method for isolation of MVs by ultracentrifugation method*, E. coli* were grown by inoculating from single colony at 120 rpm up to OD reached ∼ 1.0. The cells were pelleted, and supernatant was discarded subsequently, bacterial cell was resuspended in PBS buffer. Again, the cells were centrifuged at 5000 rpm for 10 minutes at 4°C temperature. Supernatant was collected and filtered through 0.22 µM by using vacuum filtration system. Filtered solution contains MVs and were then ultracentrifuge at 100000 g for 3 hours at 4°C. The pellet obtained resuspended in 1 mL of PBS having bacterial MVs. Isotherm analysis for interaction studies of bacterial MVs with cells and reconstituted bio-mimicking model membrane systems. Lipid molecules from bacterial MVs were isolated using the Folch method ^11^. 300 µL of freshly prepared MV solution in MV buffer was dried with nitrogen gas stream and kept in vacuum chamber overnight for buffer evaporation. 300 µL chloroform was then added to the dried bacterial MVs and bath sonicated for 30 minutes at constant pulse of 1sec. Post sonication, the solution was transferred to another vial and again dried with nitrogen gas. Chloroform (320 µL) and ice-cold methanol (160 µL) were subsequently added in 2:1 v/v ratio to dried sample and incubated for 20 minutes on ice with occasional vortex mixing followed by addition of 150 µL deionized water and kept on ice for additional 10 minutes with occasional agitation. The mixture was then centrifuged at 1500 rpm for 5 minutes for removal of the upper polar phase containing salts and proteins. For additional washing, 2:1 v/v chloroform and ice-cold methanol was again added, and the above process was repeated. After removal of the upper phase, the final lower organic phase containing lipids was dried with nitrogen stream and suspended in 100 µL chloroform. It was kept at −20°C for future use. Extracted lipid estimation of bacterial MVs is shown in the supporting text ^11^.

### Size Distribution, Detection and Fluorescence Microscopy of MVs

The detection and size of purified bacterial MVs was confirmed by fluorescence microscopy and dynamic light scattering (DLS) experiments. Fluorescence imaging and fluorimetry were done by incorporating a membrane binding dye Nile red to bacterial MVs. In sample preparation for imaging and fluorimetry, 990 µL of bacterial MVs in MV buffer were incubated with 10 µL of Nile red (100 µM) in 100/1 (v/v) ratio of MV/dye for 30 minutes. For control sample to perform fluorimetry for Nile red in MV was done by taking 10 µL dye in 990 µL of MV buffer. For fluorescence imaging 10 µL of Nile red labelled bacterial MVs were dropped on coverslip to visualize MV under fluorescence microscope (Olympus, IX71) with oil immersed 63x magnification of objective. Fluorescence intensity measurements of bacterial MVs were carried out by checking fluorescence intensity of Nile red incorporated bacterial MVs in MV buffer with respect to control sample i.e., MV buffer with Nile red. Excitation and emission spectra were optimized at 530 nm and 570 nm respectively.

For size distribution of bacterial MVs, 1 mL of MV in MV buffer were filled in quartz cuvette and allowed to be equilibrated for 5 minutes. DLS was measured by Zetasizer Nano ZS (Malvern Zetasizer Nano ZS90, Netherland) with 633 nm Laser by electrophoretic mobility of bacterial MVs in solution.

### Electron Microscopy of Bacterial MVs

To visualize the morphology of bacterial MVs, we performed transmission electron microscopy (TEM). For TEM analysis of isolated MVs, we fixed the MVs with 1% glutaraldehyde for 2 hours at 4°C. Fixed samples were negatively stained with 2% Uranyl acetate followed by washing twice. The stained MVs kept in vacuum for dehydration of sample and visualized under JEOL F200 equipped with Gatan camera.

### Fluorescence Labelling of Bacterial MVs with DiO and Di-8-ANEPPS

Purified bacterial MVs were labelled with a lipophilic dye DiO (3,3’-Dioctadecyloxacarbocyanine Perchlorate) and Di-8-ANEPPS to study affinity/fusion of bacterial MVs with reconstituted membranes. Both dyes mixed to MVs in ratio of 1:100 v/v dropwise with constant stirring at 37°C for 30 minutes for incorporation of dye in bacterial MVs. To separate unlabelled dye from labelled bacterial MVs, the mixture was washed PBS buffer four times using 10kD cut-off concentrators (Merck-Millipore) and dialyzed against PBS by changing the buffer twice. Labelled bacterial MVs were kept at 4°C and can be stored up to one week for further experiments.

### Fusion of Bacterial MVs to Macrophage Cells

To check the fusion of bacterial MVs with macrophages cells, fluorescence intensity was measured. Differentiated macrophages (THP-I) were freshly cultured in RPMI media from the maintained culture. Cells were grown up to 70% of confluence. Cells were washed with 2 mL PBS (pH – 7.4). Further, cells were washed with the 2 mL of PBS buffer containing bacterial MVs to monitor binding and fusion of MV in cells and observe the internalization of MVs after 2.5 hours. For sample preparation for fluorimetry to examine fusion of MVs with cells, 100 µL of cell soup was taken, re-suspended in 900 µL of MV buffer and 10 µL of Nile red. For control only 100 µL of MV, 900 µL of MV buffer and 10 µL of Nile red was mixed. Fluorescence intensity was then measured at 530 nm of excitation spectrum and 570 emission spectra in UV-visible spectrophotometer at time t=0. The same samples were scanned for 2.5 hours with 30 minutes of interval for fusion studies.

### Reconstitution of Fluorescence Labelled Giant Unilamellar Vesicles (GUVs)

Giant unilamellar vesicles (GUVs) were prepared using polyvinyl alcohol (PVA) gel-assisted method as previously described by Weinberger *et. al.* with slight modifications ^12^. PVA-coating on the surface of glass slides were used as film for swelling of lipid film for reconstitution of GUVs. To prepare PVA coated slides, PVA was dissolved in deionized water in 5% w/w ratio of PVA/water with constant stirring at 90°C. From this PVA solution, 200-300 µL is evenly spread on the glass slide and dried in oven at 50°C for 30 minutes which makes a thin film over the surface of slide. This dried PVA-coated slides then cleaned with UV for 15 minutes to prevent dewetting of PVA coating. Now, 20 µL of Lipid solution in chloroform (1mg/mL) doped with 1 mole % Rhodamine-PE of total lipid mole fraction was smeared on a PVA-coated slide using a Hamilton syringe. Residual chloroform on the PVA-coated slide was Evaporated by keeping the slide in vacuum chamber for 2 to 3 hours. A chamber on the lipid spread PVA-coated slide was made using nitrile ring sealed by using a second coverslip and clips as sealant. This lipid film was then swelled by filling the chamber with 1 mL of 10 mM phosphate buffer saline (PBS) at pH of 7.4 for vesiculation. The buffer used for hydration as well as the buffer with the suspended MVs were adjusted for equi-osmolarity. After 30 minutes the chamber was gently agitated, and the buffer of chamber was collected in microcentrifuge tubes containing GUVs.

### Confocal Fluorescence Microscopy

Custom made open chambers were used for incubating the Rhodamine-PE-GUVs and DiO-MVs and NBD-PE-GUVs and Di-8-ANEPPS. The cover slip was rinsed with 70% ethanol and air-dried. 10 μL of the GUVs from the was added to 90 μL of equi-osmolar MV buffer and 1mg/mL bacterial MVs were injected, and samples were incubated for 15-20 mins and allowed to equilibrate and binding of MVs. Imaging was performed using a Leica TCS-SP8 confocal instrument using appropriate lasers for rhodamine-PE (DPSS-561) and DiO (argon-488). An identical laser power and gain settings were used during the course of all the experiments. The image processing was done using ImageJ.

### Preparation of Large Unilamellar Vesicles (LUVs)

Each specific type of LUV was prepared with the help of different phospholipids in total number of 120 nmoles by corresponding lipids dissolved in chloroform. The mixture of lipids was doped with 1 mole% of di-8-ANEPPS and for background correction only lipids were mixed free of dye. The prepared mixtures then dried with steady nitrogen gas stream and kept in vacuum for 3 hours for complete evaporation of residual solvent from the lipid mixtures. Thereafter, lipids were re-suspended in 500 µL of MV buffer (pH-7.4) for hydration of lipids then suspension was heated in water bath for 15 minutes at 45 °C. After heating, samples were vortexed for 5 minutes and sonicated for 2 minutes at 0.9 sec pulse rate and 100% amplitude to produce LUVs of roughly 500 nm diameter which was confirmed by dynamic light scattering (DLS).

### Anisotropy and Dipole Potential Measurement of Reconstituted Membranes

Anisotropy and dipole potential measurements of specific lipid membranes were done by fluorescence spectroscopy. For these studied, reconstituted LUVs with specific composition of phospholipids were used and incubated with bacterial MVs. For each sample preparation, 200 µM of LUVs without bacterial MVs was taken as control whereas the same sample treated with 0.2 mg/mL and 0.6 mg/mL of bacterial MVs were the test samples. Now, these samples were incubated for 1 hour at 25°C in dark before fluorometric quantification i.e., anisotropy and dipole potential measurements. All measurements were done in 500 µL of quartz cuvette by using LS55 PerkinElmer spectrophotometer. For dipole potential study, excitation spectra were obtained by scanning the sample with 5 nm slit bandpass at 240 nm/s scan speed and the emission wavelength was fixed at 670 nm to avoid membrane fluidity effects. Background was corrected by subtracting the intensities of test samples containing di-8-ANEPPS with sample without the dye. Dipole potential was calculated by taking ratio (R), of emission intensities at 670 nm excited at 420 nm and 520 nm respectively for each sample. All values of anisotropy were calculated automatically by the instrument by using the following equation (1)^13^:

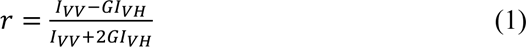

Where, I*_VH_* and I*_VV_* are the measured fluorescence intensities with excitation polarizer oriented vertically and emission polarizer oriented horizontally and vertically, respectively. G (= I*_HV_*/I*_HH_*) is the grating correction factor and is the ratio of the efficiencies of the detection system for vertically and horizontally polarized light. All experiments were conducted with multiple independent sets for each lipid.

### Lipid Mixing Experiment performed with labeled liposomes

Fusion between LUVs of outer leaflet model membrane (OLMM-RhodPE) composed of (DOPC 35.7 mole%, DOPE 5.5 mole %, SM 19.2 mole%, and Cholesterol 31.3 mole%) labeled with NBD-PE/Rh-PE (0.8 mol% each) and unlabeled bacterial MVs were monitored by the NBD fluorescence. The kinetics of lipid transfer from LUVs to MVs through FRET dilution was measured as a function of time. The lipid transfer during vesicle fusion was monitored using the change in FRET efficiency between NBD-PE (donor) and Rh-PE (acceptor). The emission intensities of the donor were monitored in 96 well Clariostar micro plate reader with excitation and the emission wavelength monochromator set to 465 nm and 530 nm, respectively. Each experiment was repeated at least three times. The extent of lipid mixing was calculated using following equation (2):

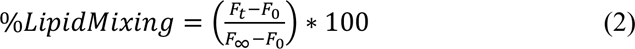

where ‘*F_0_*’ is the initial fluorescence, ‘*F_t_*’ is the fluorescence after t minutes of incubation, and *F_∞_* is the value of fluorescence after addition of TX-100 (1%) to dissolve the liposome, which is considered as the maximum disperse of the probe ^14^.

### Langmuir Blodgett Monolayer Preparation and Isotherm Analysis

Langmuir monolayer films were prepared using a microprocessor-controlled Teflon molded LB trough (Apex Instruments, India) having inner working dimension of 305 mm X 105 mm. The rectangular trough was equipped with two movable Teflon bars which provided symmetrical compression speed and the surface pressure being measured with an accuracy of ±0.05 mN/m by a Wilhelmy plate (filter paper of dimensions 10 x 25 mm^2^) connected to a highly sensitive electronic balance system. The entire system was set inside a transparent glove box. The trough was cleaned subsequently with methanol, ethanol and ultrapure water to make it dust free and utmost care was taken to avoid internal and external vibration during the experiments. MV buffer was used as the subphase whose temperature was maintained at 25 °C. Prior to each isotherm run, the interface cleaning operation was performed by compressing the two bars to a maximum and checking the surface pressure. Lipid solution at 1 mg/ml concentration was spread drop by drop on the air/ buffer interface until a surface pressure of 2-3 mN/m was reached and waited for 15-20 minutes for the chloroform to evaporate off and π to come down to 0 mN/m. 10 µL MV solution (2 µM) was injected into the subphase prior to monolayer compression. The subphase was constantly agitated using a magnetic bead stirrer at the bottom of the trough for uniform distribution of bacterial MVs. π-A Isotherm run was started by compressing the monolayer at a constant speed of 8 mm/min. Isotherms were recorded until collapse pressure π_c_ was reached. Thorough cleaning of the trough was followed each run. Each isotherm was repeated independently to ensure consistency. Isotherm data were used to calculate compressibility modulus (C_s_^-1^), excess Gibb’s free energy of mixing (DG*_Excess_*) and bending force (ΔF_b_).

### Statistical Analysis

Data were analyzed by one-way analysis of variance (ANOVA) using Origin 8.5 software. *p-values* < 0.5 were considered were considered statistically significant. All values were reported as mean ± SE from appropriate sample size and three independent experiments as indicated and where (***) depicts p<0.001, (**) p<0.01 and (*) p<0.05.

## Results

### Bacterial membrane vesicle (MV) fusion increases the fluidity and dipole potential of host lipid membrane

Bacterial membrane vesicles have been reported to enter host cells via multiple routes such as endocytosis, micropinocytosis and fusion in different cell types ^15^. Here, we sought to elucidate the lipid specificity and modulation of host membrane properties by bacterial MV interaction which remains central to all routes of entry yet remains unexplored. We used *E. coli* as a model Gram negative bacterium for the isolation of bacterial MVs and set out to systematically investigate the interaction of bacterial MVs with host membranes (***Fig. 1A***). Bacterial MVs were isolated by means of chemically induced vesiculation, which in comparison to the usual ultracentrifugation-based isolation, yield a relatively homogenous size of MVs. The characterization of the size heterogeneity of bacterial MVs was carried out by electron microscopy, fluorescence-based pixel size estimate and dynamic light scattering. Electron microscopy of MVs (***Fig. 1B, Fig S1A-S1D***) showed that MVs isolated by ultracentrifugation-based method had a broad size range of ∼ 50-200 nm (***Fig. 1C***), unlike, chemically induced vesiculation method that yield MVs with diameter of ∼ 50-60 nm (***Fig. 1C, Fig S1***). The same was found to be ∼ 150 nm by dynamic light scattering (***Fig. 1D, Fig S1E***). Further, we labelled bacterial MVs with fluorescent lipophilic dyes - such as 3,3′-dioctadecyloxacarbocyanine perchlorate (DiO) (***Fig. 1E***), Nile red and Di-8- ANEPPS to monitor binding and visualization of MV fusion by confocal microscopy as they are known to fluoresces only under lipid-rich environment ^16^.

**Figure 1.**
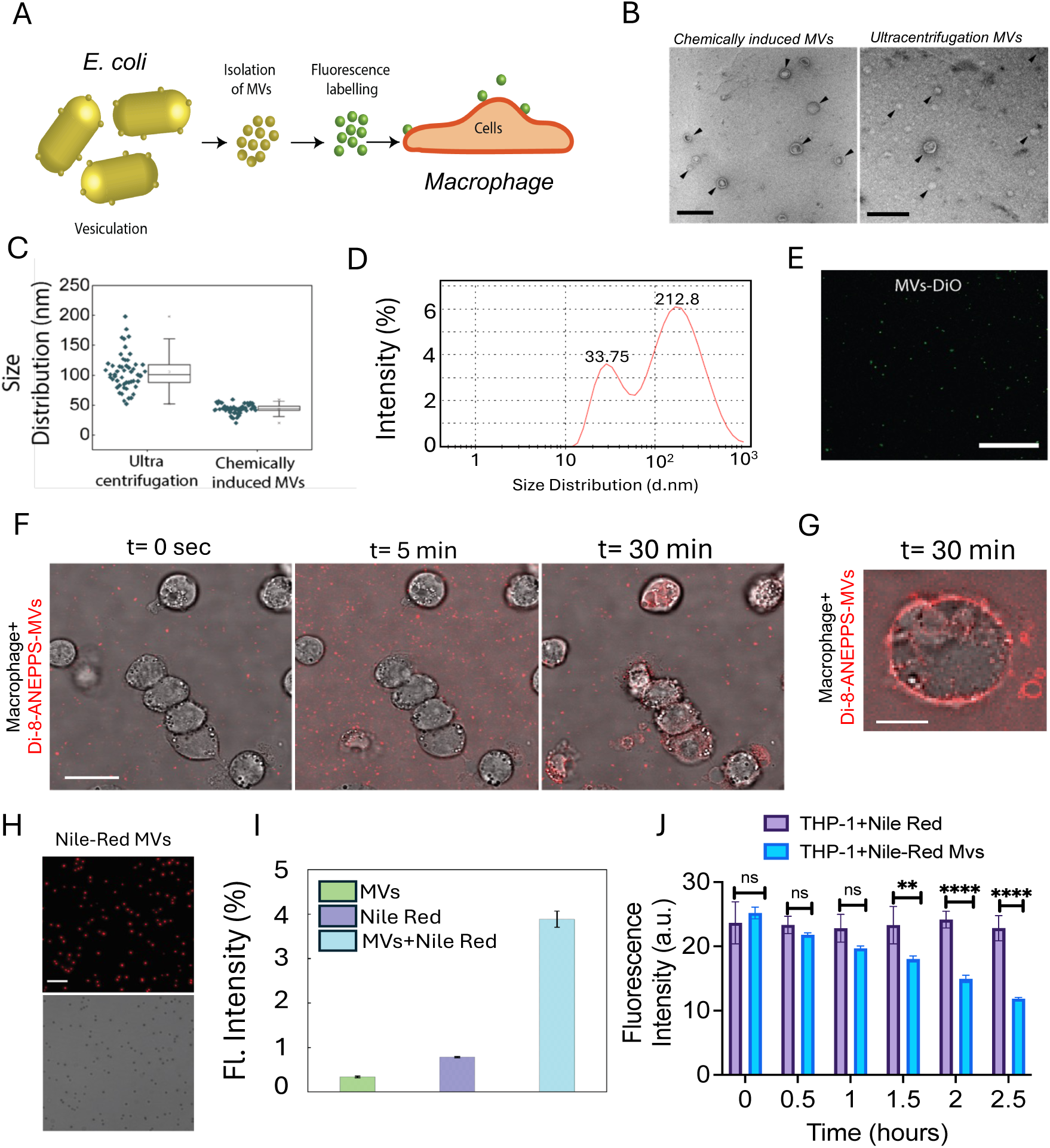
Characterization of bacterial Membrane Vesicles (MVs) isolated from *Escherichia coli* and internalization of bacterial MVs in macrophages. A) Schematic summarizing the methodology used to isolate, fluorescently label and study the MV-host membrane interaction. **B)** TEM micrograph of MVs isolated through chemical vesiculating and ultracentrifugation method (scale bar – 500 nm). **C)** Box plot for size distribution of MVs obtained through TEM micrographs. **D)** Plot showing size distribution of MVs by DLS method. **E)** Confocal image of DiO labelled MV suspension (Scale bar −10 μm). **F)** Time-course confocal sections showing fusion and internalization of Di-8-ANEPPS labelled MVs (red fluorescence) in differentiated macrophages-THP-I cells. Scale bar is 25 µm. **G)** Image from a different plane of THP-I cells with Di-8-ANEPPS after 30 minutes of incubation. Scale bar is 20 µm. ∼10-12 cells were visualized under the microscope. **H)** Fluorescence image of MVs labelled with Nile red dye (scale bar is 10 μm). **I)** Bar plot depicting fluorescence intensity of MVs (without Nile Red), Nile red only, and MVs labelled with Nile Red. **J)** Histogram showing fluorescent intensity measurements of Nile red incubated with macrophages and Nile Red-MVs incubated to differentiated macrophages with increasing at different time-points to confirm fusion of MVs to host cell. Values are mean intensity ± S.E. Significance was determined by one-way ANOVA with indicating significant difference; ns= non-significant, **p < 0.01, ***p <0.001, ****p <0.0001.

We then confirmed the interaction of bacterial MVs by human macrophages using differentiated THP-1 cells wherein significant internalization of the Di-8-ANEPPS labelled MV can be seen inside the macrophage after 30 minutes of incubation (***Fig. 1F-1G***) ^9, 17^. Additionally, the labelled bacterial MVs (devoid of any free unbound dye) were incubated with the THP-1 cells in cell media for a period of about 2.5 hours at 37°C to monitor the changes in the bacterial MV fluorescence arising in the cell media. Measurement of fluorescence intensity of a fixed volume of isolated cell media every 30^th^ minute revealed a gradual yet linear decrease in the fluorescence arising from the residual labelled MV (***Fig. 1I***). This methodology also provides an indirect measure of the kinetics of the bacterial MV fusion with the macrophage cells. Fluorescence bleaching can be ruled out as no significant change in fluorescence of the cell media containing MV was observed in the absence of macrophage cells (***Fig. 1J***).

Previous investigations have reported that Bacterial MVs not only associate with the host cell surfaces but also incorporate with different phospholipid liposomes in vitro^9^. Irrespective of the kind of bacterial MV, the interaction of the bacterial MVs takes place with the outer leaflet of the host lipid membrane. To this end, we reconstituted giant unilamellar vesicles (GUVs) comprising the predominant lipids of the outer leaflet of a eukaryotic cells (i.e., DOPC 35.7 mole%, DOPE 5.5 mole %, SM 19.2 mole%, and Cholesterol 31.3 mole%), in order to mimic the outer leaflet model membrane (OLMM) of host cell membrane^18^.

Significant variations were compared through one-sided ANOVA of OLMM treated with 0.2 mg/mL and 0.6 mg/mL of Bacterial MVs with respect to control, where, (***) depicts p<0.001(**) p<0.01 and (*) p<0.05, data point with ‘ns’ mark are not significant. All representative isotherms are the mean of independently performed experiments. For isotherms and compressibility modulus curves the estimated standard error for data points of surface pressure (π) with respect to area per molecule (A) was approximately ±0.05 mN/m.

To corroborate the MVs binding to OLMM, we labeled GUVs with Rhodamine-PE (emission wavelength, 576 nm) and bacterial MVs with DiO (emission wavelength, 501 nm). The binding of DiO labelled bacterial MVs was observed with the outer leaflet model membranes (OLMM) (***Fig. 2B***). To rule out the passive membrane incorporation equilibrium of DiO, we labelled the large unilamellar vesicle of DOPC with DiO as a control and incubated with outer leaflet model membrane (OLMM).

**Figure 2.**
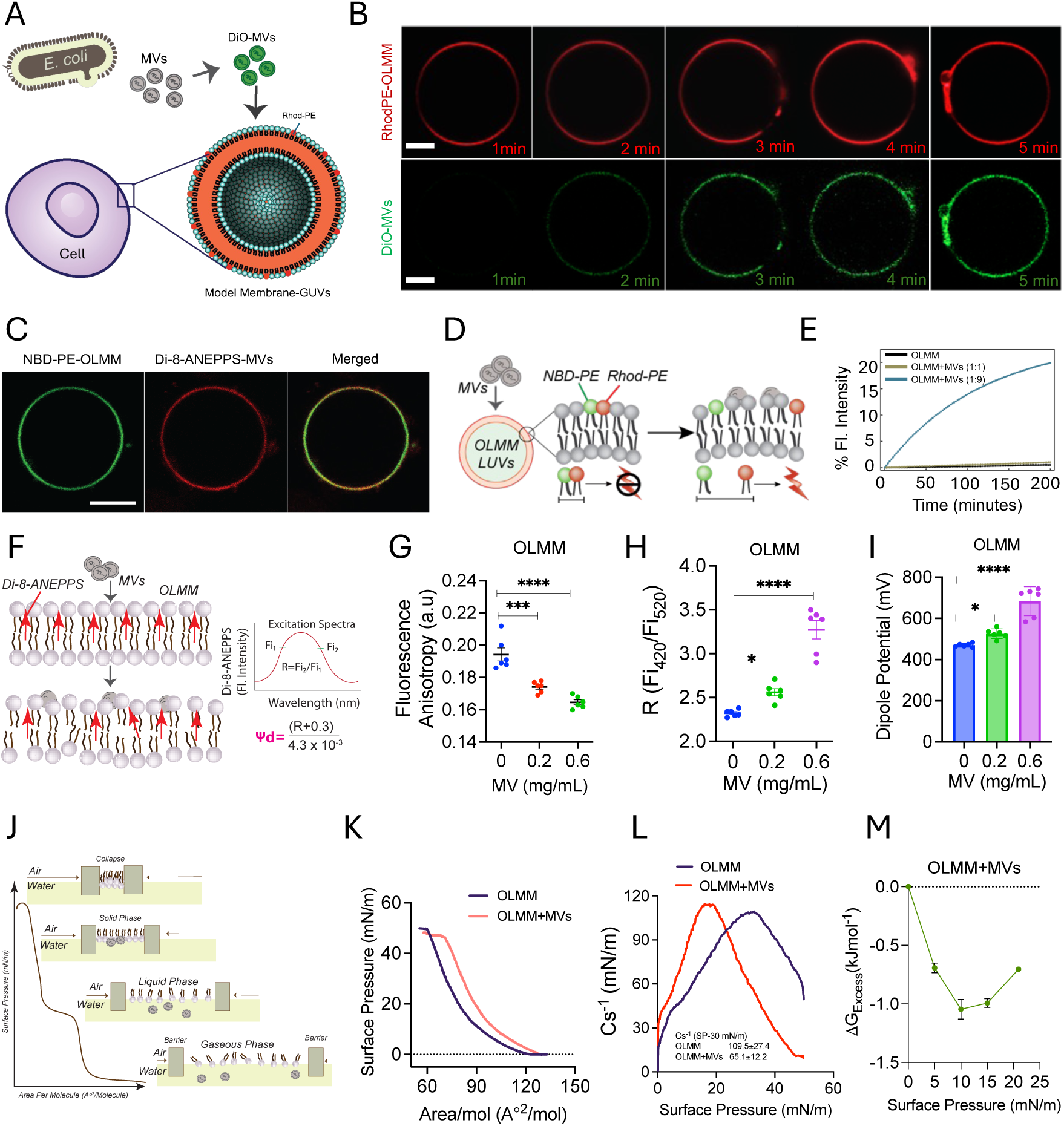
Bacterial membrane vesicles (MVs) binds to host model membrane and alters biophysical parameters of host lipid membrane. A) Model depicting the approach to study binding of fluorescently labelled-MVs with host lipid model membrane (Giant Unilamelar Vesicles). **B)** Confocal images of reconstituted GUVs of outer leaflet model membrane (OLMM-RhodPE) composed of (DOPC 35.7 mole%, DOPE 5.5 mole %, SM 19.2 mole%, and Cholesterol 31.3 mole%) doped with 1 % Rhodamine PE (*red channel*) mimicking outer leaflet of eukaryotic cell membrane. Bacterial MVs labelled with DiO binding to the GUVs (green channel) distributed homogeneously over the equatorial plane of GUVs along with merged image of both red and green channels showing a co-localization. **C)** Confocal sections showing the distribution of Di-8-ANEPPS-MVs (red fluorescence over the host lipid membrane NBD-PE-OLMM (green fluorescence).. Fluorescence imaging was done thrice independently and at least 30-35 GUVs were observed under microscope (scale bar −10µm). **D)** Illustration is showing experimental design of FRET assay for MVs-OLMM mixing. LUVs (outer leaflet) doped with quenching dye pair-NBD-PE and Rhodamine-PE and incuabted with MVs and fusion of MVs would increase Intermolecular distance of both dye and fluorescence intensity will increase. **E)** Lipid mixing kinetics of labeled LUVs with bacterial MVs at 37 °C. An increase in NBD fluorescence implies fusion between LUVs and MVs as compared to control. All measurements were carried out in PBS buffer, pH 7.4. The values represent means of at least three independent experiments performed in triplicate ± S.E. **F)** Schematic diagram depicting intercalating of charge-shift probe Di-8-ANEPPS to the cell membrane and excitation spectra showing changes in the orientation of the probe in the membrane bilayer. **G) and H)** Fluorescence intensity ratio at 420/520 nm of wavelength and dipole potential of the outer leaflet model membrane incubated with increasing concentration of Bacterial MVs, increased at high concentration of Bacterial MVs. **I)** Fluorescence anisotropy of OLMM decreases with increasing concentration of bacterial MVs correlating with the increased fluidity of host model membrane. **J)** Model showing Langmuir-Blodgett set up for lipid monolayer formation and phases of monolayer with compression. MVs injected in water subphase to observe effect on OLMM. **K)** Surface Pressure (π) -Mean molecular area (A) isotherm of OLMM monolayer (black curve) and OLMM incubated with bacterial MVs (red curve) showing right shift. All isotherms were compressed with constant speed of 8 mm/minute at 25 °C temperature. **L)** Compressibility modulus (C_s_^-1^) with respect to surface pressure of OLMM monolayer (black curve) and OLMM with MV (red curve) showing decreased C_s_^-1^ on the air-water interface. **M)** Excess Gibb’s free energy of mixing between OLMM and bacterial MVs shows an ideal mixing and decrease with increasing surface pressure calculated from Langmuir isotherms. All fluorescence spectroscopic data are shown as mean ± S.E. from three independent experiments.

No binding or fusion of DiO-DOPC to host model membrane was observed as monitored by confocal microscopy (***Fig. S2***). On the contrary, we investigated the MVs binding by labelling the MVs with membrane incorporating dye – Di-8-ANEPPS and GUV comprised of the outer leaflet lipids of the host membrane by NBD-PE. To rule out bleed-through due to wide spectra of Di-8-ANEPPS, we limit the emission bandwidth of Di-8-ANEPPS to 10% of NBD-PE excitation, which would minimize the spectral overlap of both dyes (***Fig S8***). The homogenous distribution of MVs over the equatorial plain of the of NBD-PE-OLMM GUV suggests significant interaction of MVs with host lipid membrane (***Fig. 2C***). Furthermore, The binding of bacterial MVs to the host membrane mimic comprising outer leaflet lipids as well as THP1 cells appeared to take place within 30 minutes as reported earlier in different cell lines ^19^.

The fusion of bacterial MVs with membranes was further established using the Fluorescence resonance energy transfer (FRET) between labeled LUVs and bacterial MVs. We used the fluorescent FRET pair NBD-PE and Rh-PE, where enhancement in donor (NBD-PE) fluorescence intensity shows lipid mixing taking place between the two vesicles indicating fusion (***Fig 2D***). To examine the fusion event, we have taken two different ratios of probed LUVs (i.e., containing the FRET pair) with MVs to monitor the maximum fusion at 37°C and at neutral pH 7.4. Fusion between probed LUVs and unprobed LUVs was monitored as a control which showed no significant change in donor fluorescence even after incubation for 3-4 hours. The probed to unprobed (LUV:MV, 1:1) membrane system showed only ∼1% maximum lipid mixing. A ratio of 1:9 has previously been reported and widely used for such membrane fusion assays^20^ and upon taking probed LUV and unprobed MVs in 1:9 ratio, a lipid mixing of ∼20% was observed as shown in ***Fig. 2E***. The time course of lipid mixing is much significant when MV concentration is higher, which indicates probed vesicles get fused with bacterial MVs. This is expected as the probability of interaction of MVs with the host membrane would increase with the number of MVs secreted and the amount available in the vicinity of the host membrane for random collision and interaction

We next sought to investigate whether the interaction of the bacterial MVs with the host membrane results in changes in the fluidity and dipole potential (Ψ_d_) of the host membrane (***Fig. 2F***). The changes in membrane fluidity were quantified by reconstituting a potentiometric styryl membrane probe di-8-ANEPPS (Pyridinium, 4-[2-[6-(dioctylamino)-2-naphthalenyl]ethenyl]-1-(3-sulfopropyl)), with membrane vesicles, whose fluorescence anisotropy (a measure of rotational relaxation time) responds to the changes in the dynamics of the membrane environment (***Fig. 2F***). The fluorescence anisotropy of the host outer leaflet model membrane was found to decrease by ∼ 9 % upon MV interaction suggesting increased fluidization of the host membrane (***Fig. 2G***). Likewise, the probe di-8-ANEPPS, is known to undergo shift in the fluorescence excitation spectrum in linear response to dipole potential changes in the lipid environment, which in turn, is dependent on the electric field in the membrane environment ^13, 21, 22^. The dipole potential of the host outer leaflet model membrane was found to increase by ∼ 67 % suggesting strong local remodeling upon the interaction of MV (***Fig. 2H-2I***). The dipole potential originates from the non-random arrangement of molecular dipoles (i.e., water, carbonyl-groups and lipid head-groups) at the membrane interface ^22^. Further, the surface zeta potential of the bacterial MVs was found to be ∼ −9 mV and that of the outer leaflet model membrane lipid components was also found to be negative, however, of higher magnitude (−20mV to −25mV) as compared to bacterial MVs (Table S1).

### Bacterial MV fusion increases the Compressibility and facilitates bending of host model membrane

After establishing the internalization of MVs in cells as well as binding and fusion with host model membranes, estimating the accompanied changes in the fluidity and dipole potential of the host membranes, we further sought to quantify the changes in the Compressibility of the host membrane. Langmuir monolayer surface area-pressure isotherms are a powerful means that enable quantification of the thermodynamic aspects of the MV interaction with the host model membranes and the associated changes in physicochemical properties of the host membranes^23^. We first investigated the surface pressure-area (π-A) isotherms (***Fig. 2J***) for the outer leaflet model membrane (OLMM). OLMM being a complex multi-component lipid mixture only showed liquid-expanded (L_e_) and -condensed (L_c_) behaviour, without any significantly evident 2D phase transition (***Fig. 2K***). Marked increase in the area per molecule with increasing surface pressure was observed, in the presence of MV in the sub-phase suggesting significant interaction and fusion between the monolayer membrane and bacterial MVs (***Fig. 2K***). To further corroborate our conclusions, the changes in the host model membrane surface compressional modulus, C_s_^-1^ (i.e., reciprocal of surface compressibility or Compressibility, C_s_), were extracted using the slopes obtained from (π-A) isotherms. Compressibility modulus, C_s_^-1^, is related to the change in surface pressure and area per molecule, as defined in equation (3) -

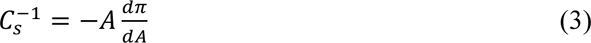

and is known to be very sensitive to subtle changes in lipid structure during lateral interaction and thus, provides molecular insights into lateral packing Compressibility of the host membrane ^24^. Upon the fusion of bacterial MVs, the monolayer compressibility modulus was found to have decreased by ∼ 44 mN/m (***Fig. 2L***), particularly nearing a surface pressure of ∼30 mN/m which has been established to possess identical chemical potential and molecular area to that of bilayers in equilibrium ^25^. Further, the excess Gibbs free energy of mixing of bacterial MVs with OLMM was calculated at certain surface pressures by integrating the excess area over surface pressure, and denoted by the equation (4) ^26^ -

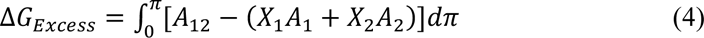

where, A_12_ is the mean molecular area occupied by the mixed monolayer in the presence of MV, A_1_ and A_2_ are the mean molecular areas occupied by monolayer and bacterial MVs respectively, and X_1_ and X_2_ are the molar fractions of the monolayer component and bacterial MVs respectively. ΔG*_Excess_* showed a decreasing trend with increasing surface pressure suggesting the mixing of MV with monolayer lipids to be more favourable till a surface pressure at 10 mN/m whereas slight increase was observed at 15 mN/m and 20mN/m (***Fig. 2M***), as the maximum crowding of bacterial MVs approached as reflected in the (***Fig. S4***). The difference of the collapse pressures (π*_c_*) of monolayer membranes in the presence and absence of MV depicts the change in lateral bending force (ΔF_b_) acting on the monolayer during the fusion which was calculated using the equation (5):

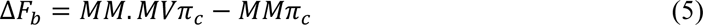

where, *MM.MVπ_c_* is the collapse pressure of model membrane with bacterial MVs in subphase and *MMπ_c_* being without bacterial MVs. Negative values of (ΔF_b_) suggest bending of monolayer ^27^. The (ΔF_b_) was found to be ∼ −1.75 mN/m suggesting the feasibility of bending upon fusion of bacterial MVs (***Table S2***). In addition, we also examined the π*-A* isotherms of the MV components (i.e., lipopolysaccharides, phospholipids devoid of any proteins) extracted from bacterial MVs (see methods for biphasic solvent extraction for lipids from MVs). A clear gaseous phase was observed in the beginning followed by L_e_ phase and a short L_c_ phase just before the collapse at around 21mN/m (***Fig. S3A***). The compressibility modulus showed a sharp rise till a surface pressure of 2-3mN/m from which it became constant at a value of 16mN/m. Both lower collapse pressure and constant compressibility modulus suggest a significantly rigid behaviour resisting compression (***Fig. S3B***).

### Role of host membrane lipid head group in bacterial MV interaction

We next questioned if there is any lipid specific preferential interaction of the OMV with the host cell membrane outer leaflet that majorly contributes towards interaction/fusion of MVs. To probe this, we reconstituted liposomes composed of the major phospholipids present in the outer leaflet of the host cell membrane i.e., DOPC, DOPG, PI and BSM. We analyzed the binding and colocalization of MVs with lipid membranes i.e. DOPC, DOPG and PI, to visualize lipid-MVs interaction (***Fig. 3A***). Significant binding of MVs to all the above lipid membranes was observed as evident from the homogenous DiO signal (green) arising from the equatorial plane of giant unilamellar vesicels (GUVs) colocalizing with Rhodamine PE signal (red) (***Fig. 3A***). After confirming the interaction of MVs with DOPC, DOPG and PI lipids, we further quantified the modulation of dipole potential and fluidity changes in these lipid membranes. We then investigated the role of lipid head group on the MV induced dipole potential changes in the host membrane. We observed that membranes composed of DOPC (***Fig. 3B-3C***), DOPG (***Fig. 3D-3E)*** and PI (***Fig. 3F-3G)*** undergo significant changes in their dipole potential (i.e., 21 %, 15 % and 9 % respectively) (***Fig. 3B-3G***) and membrane composed of BSM did not undergo any significant change in the measured dipole potential (***Fig S4B***). On the contrary, we found that DOPC, DOPG, BSM and PI membranes, all undergo significant changes in the membrane fluidity upon the interaction of MVs. Interestingly, the fluorescence anisotropy of the dye was found to decrease in case of DOPC, DOPG and Liver PI by ∼ 66 %, 45 % and 45 % respectively, suggesting enhanced fluidization (***Fig 3H-3J***). On the contrary, the membrane composed of BSM showed an increase in the fluorescence anisotropy of the dye by 7 %, suggesting the membrane becoming more rigid (***Fig. S4A***). We then sought to investigate the modulation of membrane elasticity upon MV fusion and mixing with regard to various lipid head-groups. We observed, MV addition caused decrease in area per molecule in case of DOPC (***Fig. 3K***), whereas only slight change in PI (***Fig. 3M***), which remained almost constant with increasing pressure. Coexistence of Lc+Le phase seemed to be very mild in DOPC curve with no clear occurrence in the other two lipids. On the contrary, an increase in the molecular area of DOPG in the presence of MV was observed as evident in the rise in surface pressure (***Fig 3L***).

**Figure 3.**
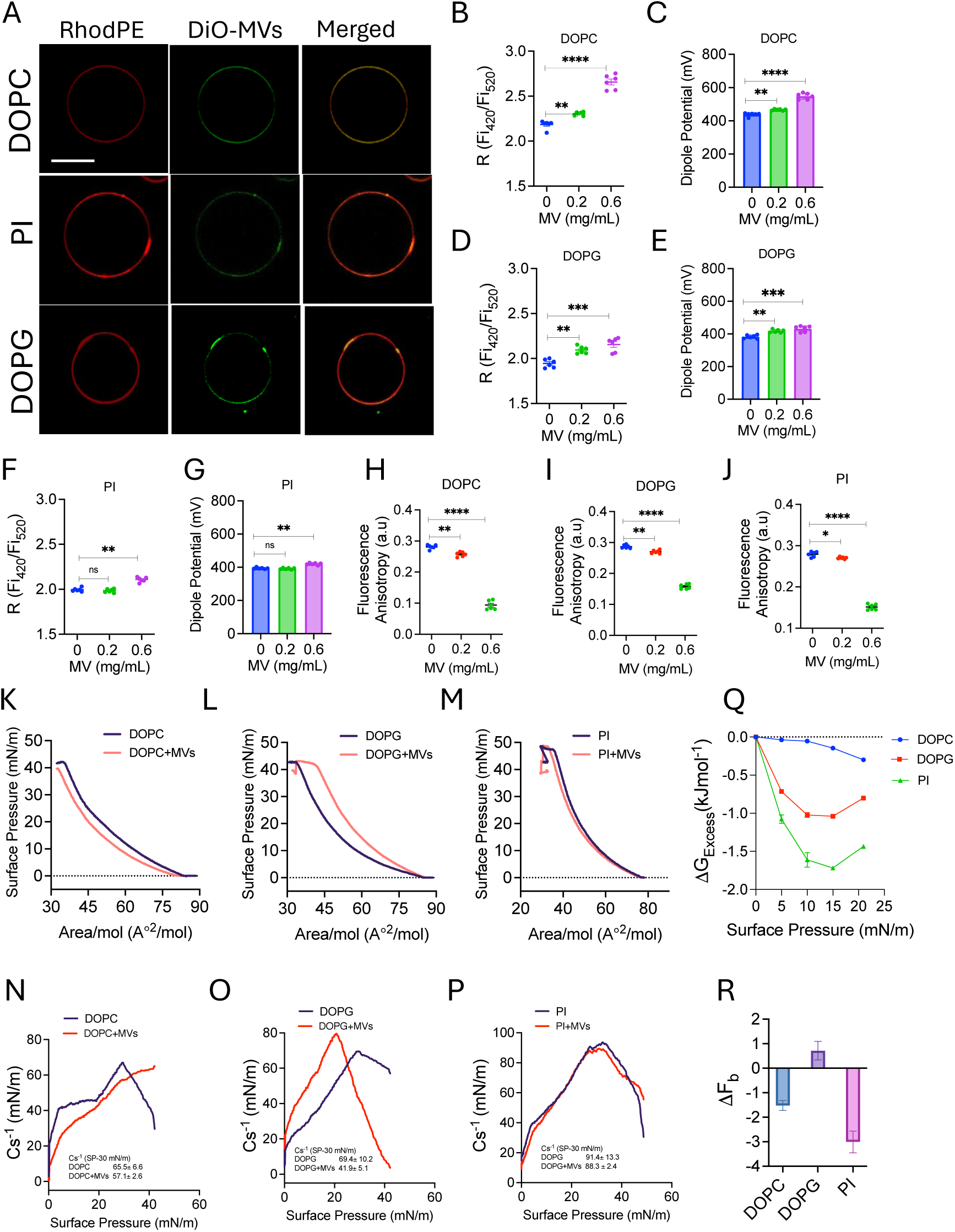
Monitoring the lipid specificity of host membrane for bacterial MVs interaction using confocal and the fluorometry of the potentiometric dye, Di-8-ANEPPS. A) Confocal image of DOPC, PI and DOPG doped with 1% Rhodamine PE (*red fluorescence)* mixed with bacterial MVs labelled with DiO (*green fluorescence)* to visualize the interaction of bacterial MVs with lipid membrane. A green fluorescence signal on the equatorial plane of membrane suggests the interaction of bacterial MVs with these phospholipids. Fluorescence ratio 420/520 nm and dipole potential (MV) for **B and C)** DOPC, **D and E)** DOPG and **F and G)** PI membranes at 0.2 mg/mL and 0.6 mg/mL concentration of bacterial MVs with respect to control model membranes. Membrane anisotropy of **H)** DOPC, **I)** DOPG and **J)** PI with increasing concentration of bacterial MVs i.e., 0 mg/mL, 0.2 mg/mL and 0.6 mg/mL. Measurements for calculation of dipole potential were done at measured intensity at excitation wavelength set at 420 nm and 520 nm respectively. All fluorescence spectroscopic data are shown as mean ± S.E. from three independent experiments. Surface Pressure (π)-mean molecular area (A) isotherms of – **K)** DOPC, **L)** DOPG and **M)** PI monolayers in presence of MVs. Changes in comp ressibility modulus of **N)** DOPC, **O)** DOPG and **P)** PI membrane derived from isotherm of respective model membrane-MVs interactions. **Q)** Variation of excess Gibb’s free energy of mixing of bacterial MVs with single monolayers (DOPC, DOPG and PI) as a function of increasing surface pressures 0 mN/m, 5 mN/m, 10 mN/m, 15 mN/m and 20 mN/m. **Q)** Bending force (ΔF_b_) histogram of DOPC, DOPG and PI, monolayers at the interface in presence of bacterial MVs. All isotherms were recorded with constant barrier compression speed of 8 mm/min at 25°C. All representative isotherms are the mean of independently performed experiments conducted thrice. For isotherms and compressibility modulus curves the estimated standard error for data points of surface pressure (π) with respect to area per molecule (A) was approximately ±0.05 mN/m.

A reduction of 1.52 mN/m and 3.01 mN/m in the collapse pressure of DOPC and PI membrane was observed, though DOPG membrane showed a little rise of 0.72 mN/m (***Fig 3K-3L***). The negative values of the change in bending force (Δ𝐹_1_) suggest bending of monolayer ^27^ by the interacting MV as observed in case of both DOPC and PI lipid membranes (***Fig. 3R***).

Further, OMV interaction/fusion induced a reduction in monolayer compressibility modulus (C_s_^-1^) by around ∼ 8 mN/m, ∼ 28 mN/m and ∼ 2 mN/m, in DOPC, DOPG and PI lipid membranes, respectively. (***Fig. 3N-3P***). The reduction in C_s_^-1^ in the presence of MV suggests a rise in the membrane monolayer compressibility. We next wanted to see if the mixing of MV with each lipid monolayer is thermodynamically favourable or not. A decreasing negative value trend in ΔG_Excess_ was observed (***Fig. 3Q***) with respect to increasing surface pressure till 15 mN/m in DOPC and PI and 10 mN/m in DOPG after which the curve rose. Upon the mixing of MV, DOPG membrane monolayer showed the lowest free energy change of ∼ −1.05 kJ/mol and highest being ∼ −1.7 kJ/mol for PI indicating more favourable mixing with PI monolayer.

### Role of lipid acyl chain in OMV mediated modulation of membrane fluidity, dipole potential and elasticity

The lipid acyl chain length is known to determine the membrane bilayer thickness as well as contributes to the degree of fluidity of the membranes ^28^. Therefore, we then sought to investigate the role of lipid acyl chain (i.e., C12, C14, C16, C18) of host membrane in the MV mediated changes in the overall host membrane fluidity and dipole potential. The overall fluorescence anisotropy of the dye was found to decrease by ∼ 30 % in DPPC suggesting increased fluidization (***Fig. 4C***), and increase by ∼ 28 %, ∼ 43 % and ∼ 27 % in DLPC, DMPC and DSPC respectively, suggesting decrease in fluidity (***Fig. 4A, 4B and 4D***). The contrasting observation is noteworthy, as it suggests, that MV interaction seems to enhance the fluidity of a relatively less fluid membrane (such as DPPC, C16) and decrease the fluidity of relatively more fluid membranes (such as C12,14). The observed fluctuations in the fluorescence anisotropy with increasing concentrations of the MV, could hint at a dynamic interaction during the crowding/interaction of the MV. Further, the dipole potential was found to increase in all cases (DMPC, DPPC and DSPC) (***Fig. 4E and 4I, 4G and 4K, 4H and 4L***) but decrease in DLPC membranes upon MV interaction (***Fig. 4F and 4J***). On the contrary, MV interaction with DLPC membranes resulted in less significant change in the dipole potential. In case of DMPC (***Fig. 4F and 4J***) and DSPC (***Fig. 4H and 4L***), the dipole potential was found to increase by ∼ 30% and ∼ 15% respectively, at highest concentration of MVs as compared to control which confirms interaction of bacterial MVs with DMPC and DSPC membrane. Likewise, DPPC membrane shows significant increase in dipole potential upon interaction (***Fig. 4G and 4K***).

**Figure 4.**
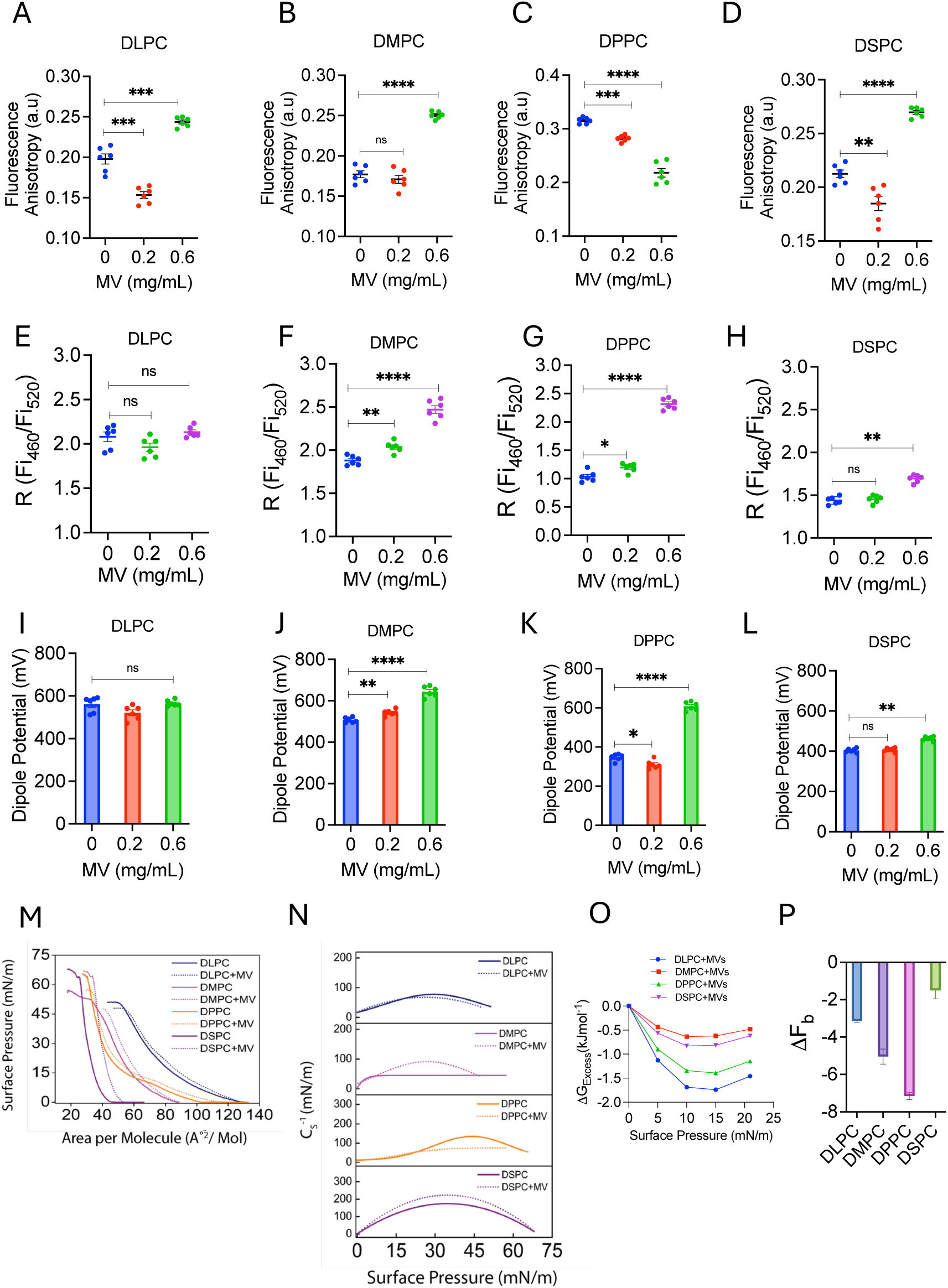
Variation in biophysical properties of phospholipid membranes with increasing bilayer thickness as a function of acyl chain length, treated with MVs. Fluorescence anisotropy of individual lipid model membranes **A)** DLPC, **B)** DMPC, **C)** DPPC, and **D)** DSPC incubated with increasing concentration of MVs. Changes in fluorescence ratio and dipole potential of **E and F)** DLPC, **G and H)** DMPC, **I and J)** DPPC, and **K and L)** DSPC membrane with respect to increasing concentrations of MVs i.e., 0 mg/mL, 0.2 mg/mL and 0.6 mg/mL. All fluorescence spectroscopic data are shown as mean ± S.E. from three independent experiments. Significant variations were compared through one-sided ANOVA of membrane condition treated with MVs with respect to control (only lipid membrane without MVs), where (***) depicts p<0.001, (**) p<0.01 and (*) p<0.05, data point with no asterisk mark are non-significant. π-A isotherms of **M)** DLPC (12:0), DMPC (14:0), DPPC (16:0) and DSPC (18:0) with and without bacterial MVs obtained at 25°C**. N)** Variability in compressibility modulus plotted as a function of surface pressure acquired for DLPC (12:0), DMPC (14:0), DPPC (16:0) and DSPC (18:0). **O)** Variation of excess Gibb’s free energy of mixing of bacterial MVs with lipid monolayer as a function of increasing surface pressures 0 mN/m, 5 mN/m, 10 mN/m, 15 mN/m and 20 mN/m. **P)** Bending force (ΔF_b_) histogram of DOPC, DOPG, and PI monolayers at the interface in presence of bacterial MVs. All isotherms were recorded with constant barrier compression speed of 8 mm/min at 25°C. All representative isotherms are the mean of independently performed experiments conducted thrice. For isotherms and compressibility modulus curves the estimated standard error for data points of surface pressure (π) with respect to area per molecule (A) was approximately ±0.05 mN/m.

We then sought to question the role of lipid acyl chain length in MV interaction/fusion using monolayer of lipids at the air-water interface. This is important as under cellular conditions transient phase separated regions might differ in the membrane thickness ^28^. Analysis of surface pressure-area curves of monolayers of increasing acyl chain length, which corresponds to membrane bilayer thickness, were in good agreement with previous reports ^29^. We observed that the mean molecular area occupied by phospholipids decrease with increasing chain length in control conditions i.e. in absence of bacterial MVs, except DPPC (***Fig. 4M***), which can be explained by the effective acyl chain hydrophobic interactions^30^. In presence of bacterial MVs, increase in area per molecule was observed in each of the four membrane monolayers with rise in surface pressure (***Fig. 4M***). The observed increase in molecular area was fairly constant throughout all the phases in DLPC, DPPC and DSPC membrane monolayers (***Fig. 4M***) whereas an increasing trend was observed in DMPC till 15 mN/m (***Fig. 4M***). Decrease in collapse pressure by 3.15 mN/m and 5.04 mN/m was calculated for DLPC and DMPC. Highest decrease in collapse pressure of ∼ −7 mN/m amongst all lipids was observed for DPPC after MV addition suggests most significant bending of membrane in 16:0 acyl chain length (***Fig 4P***). This decline was least in case of DSPC (***Fig. 4P***). ***Fig. 4N*** show monolayer compressibility curves of the lipids in presence of MV with respect to control. A decline in compressibility modulus by around 10 mN/m was seen in DLPC (***Fig 4N***), whereas a decline of 35 mN/m was observed in DPPC (***Fig. 4N***). Compressibility modulus showed a dramatic increase in case of DMPC and DSPC with a value of almost 50 mN/m and 60 mN/m (***Fig. 4N***) and lowest energy change of mixing was observed in case of DMPC throughout all pressures (***Fig. 6O and Table S3***). The ΔG*_Excess_* change though followed falling trend up to a particular pressure, there was not much difference between the Gibb’s free energy curves of varying acyl chain length containing lipids (***Fig. 6O***). At 15 mN/m, the free energy change was highest in DLPC followed by DPPC suggesting ideal mixing of bacterial MVs with these membranes.

### Lipid membrane rigidity and phase separation modulate OMV fusion

Host cell membranes are multi-component lipid assembly that is a non-equilibrium homogenous system close to phase separation. The membrane environment has highly transient phase separating regions called rafts important for cellular signaling. We therefore investigated the effect of phase behavior on the interaction of the bacterial MVs. In order to probe this, we reconstituted liposomes (GUVs and LUVs) made of widely accepted ternary membrane composition (i.e., DOPC:BSM:Cholesterol) and varied the amounts of cholesterol to enable different degrees of liquid-disordered (L_d_) and liquid-ordered (L_o_) regions. This would mimic the biologically relevant highly dynamic local changes in membrane phase boundaries ^31^. First, we reconstituted GUVs composed of DOPC/BSM without any cholesterol representing a disordered (L_d_) phase in membrane to examine how the L_d_ forming membrane condition effects the interaction of MV (*liquid disordered; DOPC:BSM:Chol 5:5:0*) ^32^. Uniform fusion of bacterial MVs to the partial disordered membrane was observed. We arrived at this conclusion by monitoring all the planes of stacks as well as 3D-stack image of GUV incubated with bacterial MVs (***Fig. 5A***). The fluorescence anisotropy of the dye increased by ∼ 27% upon interaction with MV in a concentration dependent manner indicating decrease in membrane fluidity (***Fig. 5B***). Further, the membrane dipole potential was found to linearly decrease upon the interaction of MV (***Fig. 5C-5D***), all of which hint at binding of bacterial MVs that may allow some lipid mixing ^33^.

**Figure 5.**
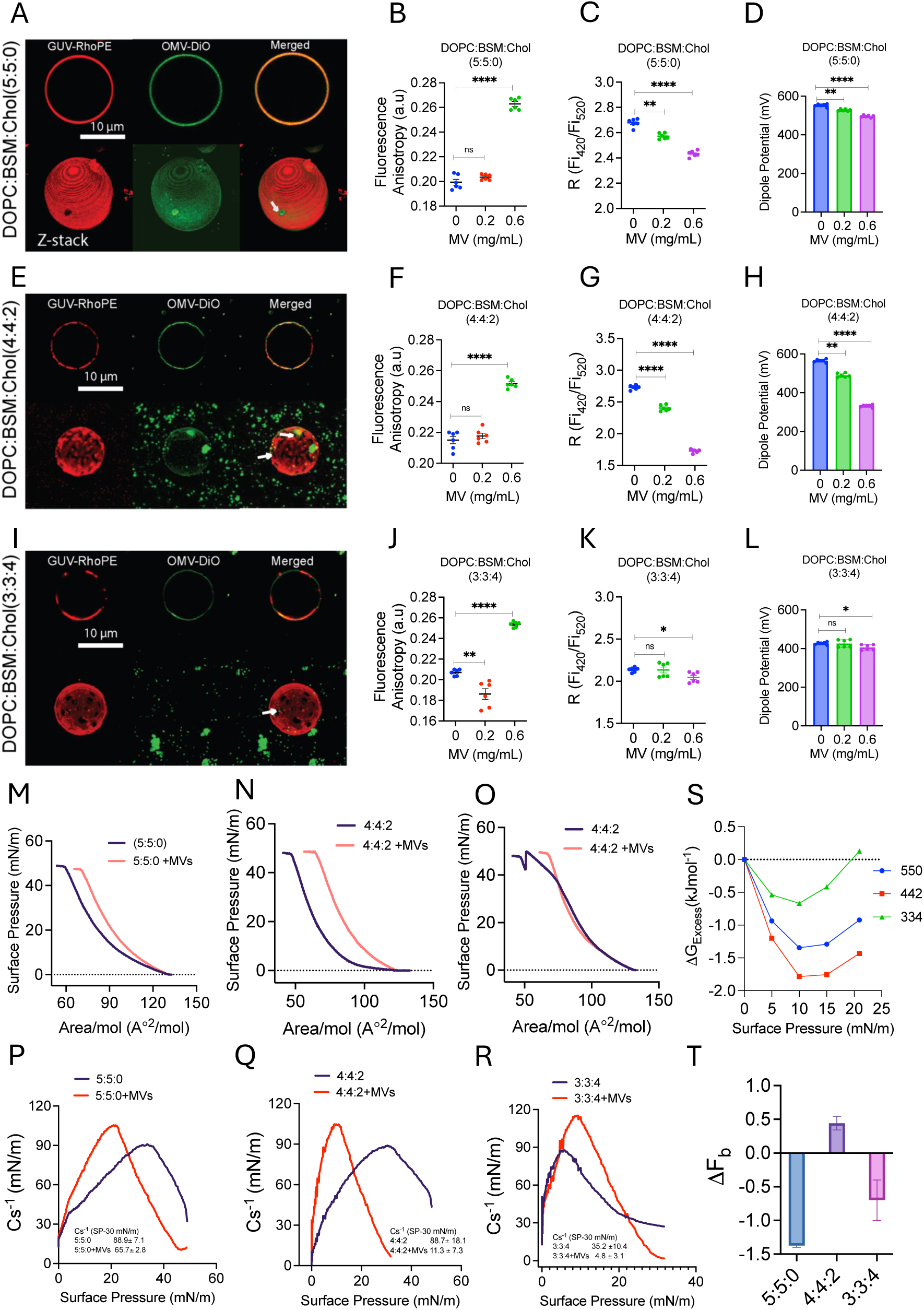
Effect of increasing concentration of cholesterol (liquid order) and phase behavior of membrane on binding, changes in membrane parameters treated with bacterial MVs. A) Fluorescence image showing GUVs composed of DOPC:BSM:Chol in the molar ratio of 5:5:0 doped with 1 mole % Rhodamine PE (*red channel)* interaction with DiO labelled bacterial MVs *(green channel)*. Bacterial MVs distribution on equatorial plane of GUVs is represented by merged image of both red and green channels along with the 3D stack of each channel. **B)** Fluorescence anisotropy of partially disordered DOPC:BSM:Chol 5:5:0 membrane induced by bacterial MVs, **C and D)** Dipole potential of partially disordered (DOPC:BSM:Chol 5:5:0) treated with increasing concentration of bacterial MVs**. E)** Multiple lipid domain can confine the binding of bacterial MVs to partially disordered DOPC:BSM:Chol 4:4:2 membrane (*left*-Rhodamine PE-(red), *centre*- DiO (green) and *right*- merged). Top panel is showing MV binding to equatorial plane of GUVs and the panel below is bacterial MVs 3D-stack of GUV-MVs mixture. **F)** Membrane fluidity variation in DOPC:BSM:Chol in molar ratio of 4:4:2 membrane treated with bacterial MVs in increasing concentration. **G and H)** Dipole potential of (DOPC:BSM:Chol) in mole ratio of 4:4:2 treated with bacterial MVs in increasing concentration. **I)** Fluorescence image showing GUVs composed of DOPC:BSM:Chol in the molar ratio of 3:3:4 correspond to liquid ordered domain (l_o_) doped with 1 mole % Rhodamine PE (*red channel)* interaction with DiO labelled MVs *(green channel)*. bacterial MVs has least binding with DOPC:BSM:Chol; 3:3:4 represented by merged image of both red and green channels along with the 3D stack of each channel. **J)** Changes in anisotropy of DOPC:BSM:Chol (3:3:4) treated with increasing concentration of bacterial MVs. **K and L)** Fluorescence ratio and dipole potential of (DOPC:BSM:Chol 3:3:4) treated with increasing concentration of Bacterial MVs. At least 30 GUVS visualized under microscope and scale bar is 10 µm. All fluorescence spectroscopic data are shown as mean ± S.E. from three independent experiments. Significant variations were compared through one-sided ANOVA of membrane condition treated with bacterial MVs with respect to control (only lipid membrane without bacterial MVs), where, (***) depicts p<0.001, (**) p<0.01 and (*) p<0.05, test data point (membrane treated with bacterial MVs) with no asterisk mark are non-significant.

We then modulated the degree of phase separation, which in turn effects the line tension at the phase separating boundaries ^34^, by reconstituting membranes having L_o_/L_d_ phase separated regions by changing the cholesterol levels (i.e, *DOPC:BSM:Chol, 4:4:2/3:3:4*). Interestingly, in case of L_o_/L_d_ phase separating membranes with moderate levels of cholesterol (*DOPC:BSM:Chol, 4:4:2*) significant binding of bacterial MVs was observed (***Fig. 5E***). The fluidity decreased in a similar manner as for the L_o_ membrane as reflected in the increase fluorescence anisotropy of the dye by ∼15 % (***Fig. 5F***). Similarly, the dipole potential was found to linearly decrease upon the interaction of MV (***Fig. 5G-5H***). Interestingly, in case of phase separating membranes comprising higher amount of cholesterol (*DOPC:BSM:Chol, 3:3:4*), having a larger L_o_ (liquid ordered) regions the MV binding was found to be relatively weak (***Fig. 5I***). Also, the fluidity was found to decrease as in earlier cases, however, no significant change in the dipole potential of the membrane (***Figs. 5J-5L***).

We then investigated the role of cholesterol in bacterial MV interaction/fusion and the subsequent mechano-elastic changes in lipid monolayer. To this end, as described earlier π*-A* isotherms of show similar trend with increase in molecular area in the presence of MV (***Fig. 5M***). A noticeable aspect observed in the monolayer with highest cholesterol content (3:3:4) in the presence of MV, was that - after surface pressure of around 7.5 mN/m, there was a left shift followed by a right shift after 35 mN/m (***Fig. 5O and 5T***). Collapse pressure reduced in both partially disordered DOPC/BSM membrane without any cholesterol and DOPC/BSM/Chol(3:3:4) while increased by ∼ 0.5 mN/m in the case of DOPC/BSM/Chol(4:4:2) membrane(***Fig. 5T, and Table S2***).

π-A isotherms of **M)** 5:5:0 (BSM:DOPC:cholesterol), **N)** 4:4:2 (BSM:DOPC: cholesterol) and **O)** 3:3:4 (BSM:DOPC:cholesterol) with and without bacterial MVs obtained at 25°C. Variability in compressibility modulus plotted as a function of surface pressure acquired for **P)** 5:5:0, **Q)** 4:4:2 and **R)** 3:3:4 from respective Langmuir isotherms. **S)** Variation of excess Gibb’s free energy of mixing of bacterial MVs with cholesterol containing membrane as a function of increasing surface pressures 0 mN/m, 5 mN/m, 10 mN/m, 15 mN/m and 20 mN/m. **(F)** Bending force (ΔF_b_) histogram of 5:5:0, 4:4:2 and 3:3:4 monolayers at the interface in presence of bacterial MVs. All isotherms were recorded with constant barrier compression speed of 8 mm/min at 25°C. All representative isotherms are the mean of independently performed experiments conducted thrice. For isotherms and compressibility modulus curves the estimated standard error for data points of surface pressure (π) with respect to area per molecule (A) was approximately ±0.05 mN/m.

## Discussion

Membrane fusion is considered one of the key mechanisms for the entry of bacterial MVs inside the host cells ^2, 15^. Using a non-pathogenic *E. coli* strain as a model Gram-negative bacterium to isolate bacterial MVs, we quantitatively investigate MVs interaction with host membrane to dissect the MV mediated modulation of host membrane fluidity, dipole potential, compressibility parameters that might change as a result of fusion and alter subsequent signalling. Fusion of bacterial MVs to host membranes is involved in transmission of virulence factors in non-phagocytic cells ^19^ as well as in reprogramming of phagosomes in phagocytic macrophage^35^. As expected, the fluorescently labelled bacterial MVs was found to undergo significant internalization by THP-1 macrophage (***Fig. 1***), in line with previous observations (11). The binding of DiO-labelled and Di-8-ANEPPS-labelled bacterial MVs to the reconstituted lipid membranes mimicking the outer-leaflet of the host cell membrane was found to be homogenous (***Fig. 2***). The enhanced FRET observed upon incubation of probed LUVs with unprobed MVs confirms fusion (***Fig. 2***). Further, observed monolayer area expansion accompanied by an ideal mixing reflected in a negative free energy change might also hint at likely fusion of MV, although adhesion not progressing into complete fusion cannot be totally ruled out. We observe that the fusion of bacterial MVs results in increased fluidization and dipole potential (***Fig. 2***). Further, it also increased the magnitude of the membrane surface potential of the host model membrane to more negative regime. Such trans-negative membrane surface potential was reported to be essential for cell-cell fusion^36^. The observed fluidization is supported by the experimentally determined increase in molecular area per lipid as well as Compressibility (as seen in decrease in the C_s_^-1^ at surface pressure at 30 mN/m) (***Fig. 2***). The surface-pressure area isotherm of the lipid fraction of MV devoid of any proteins suggests that the MV membrane is extremely rigid as evident from the low collapse pressure (***Fig. S4***). LPS, the major component of MV owing to its relatively wider cross-section area and low head-to-tail aspect ratio could be a major contributor towards fluidization as it is known to uniformly adhere and incorporate in egg-PC, and DOPE containing lipid bilayer ^37^ all of which are present in the host outer leaflet of the membrane. This is in line with recent findings that Phthiocerol dimycocerosates (DIMs), a lipid component of *Mycobacterium tuberculosis,* are transferred to and accommodated in the host cell membrane due to its conical shape ^38^.

We then wondered how would the acyl chain length effect the fusion? MV fusion to major phospholipid DOPC, DOPG and PI (***Fig. 3***) but not BSM membrane, results in enhanced membrane fluidity as well as increased dipole potential (***Fig S4***). The changes in area per lipid molecule are coupled to the degree of interfacial fusion of the interacting components ^23, 39^. The observed increase in area per lipid molecule is likely due to the formation of inverted cubic structure/aggregates at the fusion sites resulting in expulsion of some lipids ^37, 38, 40^, which, also explains the observed relatively higher negative values of G*_Excess_*. We observed that DLPC, DMPC, and DSPC membranes exhibited increased rigidity. This rigidity could be attributed to partial fusion occurring within the bilayers, resulting in a shift towards the native ordered phase of the bilayers. Interestingly, we observed that MV fusion to only DPPC membrane results in enhanced fluidization accompanied by increase in interfacial molecular dipoles unlike other lipid membrane (***Fig. 4C***). Together, the data suggest that while MV interaction is facilitated by zwitterionic head groups, however, an optimum acyl chain length is essential for effective interaction/fusion in line with previous observations ^41^.

Lipid rafts have been reported to facilitate the fusion of bacterial MVs ^8^. However, this was proved using cholesterol depleting or sequestering agents such as filipin or Methyl-ß-cyclodextrin, which cannot rule out the possibility that disruption of cholesterol on a large scale may affect protein dependent processes not limited to lipid rafts. One way to examine and correlate the exclusive dependence of MV mixing with lipid rafts is by reconstituting minimal systems of lipid raft-like regions without any protein. Bacterial MVs were found to show strongest fusion to partially disordered membranes devoid of cholesterol (DOPC:BSM:Chol, 5:5:0) accompanied by highest decrease in the fluidity (***Fig. 5***). Further, increase in the cholesterol in the membrane lead to decreased fusion of bacterial MVs to the membrane (***Fig. 5).*** The increasing cholesterol levels results in co-existing phase separated regions (L_o_ and L_d_) within the membranes (DOPC:BSM:Chol, 4:4:2 & 3:3:4), however, the number of the phase separating boundaries is higher in the former. Such differences in the phase separating boundaries would accumulate different degrees of interfacial energy in the membrane, that would facilitate the fusion of bacterial MVs in different ways, which explains the observed variation ^20, 42^.

Increase in the dipole potential of host lipid membrane by fusion of MV is likely to stimulate conformational changes in the membrane bound/inserted receptor proteins ^20, 34^. Such conformational changes in transmembrane receptor proteins have been reported to significantly alter the receptor signaling in host cells ^43^. Additionally, our study also suggests that MV interaction with lipid rafts/phase boundaries may have an important role in altering host cell signaling ^44^. *Mycobacterium ulcerans* endotoxin, Mycolactone, was shown to have potent effect on reorganization of raft-like model membranes ^45^. Although our observations are based on a non-pathogenic strain of *E. coli* as a model bacteria, it is important to note that similar observations can be expected with pathogenic *E. coIi* as they share more than 70% similarity in the protein content of the their membrane vesicles ^46^. Besides the overall changes induced by the MVs, identifying the potential proteins in the MVs and their contribution to the modulation of the host cell membrane remains an important aspect that would require extensive research.

## Author Contributions

AP and MS conceived the idea and designed the research. AP, AT carried out the characterization of bacterial MVs. DKS helped with MV characterization. AP, DK and GM performed labelling of bacterial MVs and imaging experiments. AP and TM performed monolayer experiments. AP and GM performed fluorimetry experiments. AP, AT, TM, GM and MS analyzed the data. AP, TM and MS wrote the manuscript.

## Acknowledgements

We would like to gratefully acknowledge the financial support received from the Department of Science and Technology (EMR/2017/004513) and Department of Biotechnology, Govt. of India (BT/PR/21226/MED/122/41/2016). AP, AT and TM acknowledge MHRD for the GATE fellowship. We would also like to thank Department of Atomic Energy, Govt. of India and NISER, Bhubaneshwar for financial support and infrastructure.

## References

(1) Beveridge, T. J. Structures of gram-negative cell walls and their derived membrane vesicles. Journal of bacteriology 1999, 181 (16), 4725–4733. Schwechheimer, C.; Kuehn, M. J. Outer-membrane vesicles from Gram-negative bacteria: biogenesis and functions. Nature Reviews Microbiology 2015, 13 (10), 605.

(2) Kulp, A.; Kuehn, M. J. Biological functions and biogenesis of secreted bacterial outer membrane vesicles. Annual review of microbiology 2010, 64, 163–184.

(3) Toyofuku, M.; Nomura, N.; Eberl, L. Types and origins of bacterial membrane vesicles. Nature Reviews Microbiology 2018, 1. Théry, C.; Witwer, K. W.; Aikawa, E.; Alcaraz, M. J.; Anderson, J. D.; Andriantsitohaina, R.; Antoniou, A.; Arab, T.; Archer, F.; Atkin-Smith, G. K. Minimal information for studies of extracellular vesicles 2018 (MISEV2018): a position statement of the International Society for Extracellular Vesicles and update of the MISEV2014 guidelines. Journal of extracellular vesicles 2018, 7 (1), 1535750.

(4) Koeppen, K.; Hampton, T. H.; Jarek, M.; Scharfe, M.; Gerber, S. A.; Mielcarz, D. W.; Demers, E. G.; Dolben, E. L.; Hammond, J. H.; Hogan, D. A. A novel mechanism of host-pathogen interaction through sRNA in bacterial outer membrane vesicles. PLoS pathogens 2016, 12 (6), e1005672.

(5) MacDonald, I. A.; Kuehn, M. J. Offense and defense: microbial membrane vesicles play both ways. Research in microbiology 2012, 163 (9-10), 607–618.

(6) Chattopadhyay, M. K.; Jagannadham, M. V. Vesicles-mediated resistance to antibiotics in bacteria. Frontiers in microbiology 2015, 6, 758.

(7) Kaparakis-Liaskos, M.; Ferrero, R. L. Immune modulation by bacterial outer membrane vesicles. Nature reviews Immunology 2015, 15 (6), 375.

(8) Bomberger, J. M.; MacEachran, D. P.; Coutermarsh, B. A.; Ye, S.; O’Toole, G. A.; Stanton, B. A. Long-distance delivery of bacterial virulence factors by Pseudomonas aeruginosa outer membrane vesicles. PLoS pathogens 2009, 5 (4), e1000382.

(9) Jäger, J.; Keese, S.; Roessle, M.; Steinert, M.; Schromm, A. B. Fusion of L egionella pneumophila outer membrane vesicles with eukaryotic membrane systems is a mechanism to deliver pathogen factors to host cell membranes. Cellular microbiology 2015, 17 (5), 607–620.

(10) Sezgin, E.; Kaiser, H.-J.; Baumgart, T.; Schwille, P.; Simons, K.; Levental, I. Elucidating membrane structure and protein behavior using giant plasma membrane vesicles. nature protocols 2012, 7 (6), 1042.

(11) Folch, J.; Lees, M.; Sloane Stanley, G. A simple method for the isolation and purification of total lipides from animal tissues. J biol Chem 1957, 226 (1), 497–509. Reis, A.; Rudnitskaya, A.; Blackburn, G. J.; Fauzi, N. M.; Pitt, A. R.; Spickett, C. M. A comparison of five lipid extraction solvent systems for lipidomic studies of human LDL. Journal of lipid research 2013, 54 (7), 1812–1824.

(12) Weinberger, A.; Tsai, F.-C.; Koenderink, G. H.; Schmidt, T. F.; Itri, R.; Meier, W.; Schmatko, T.; Schröder, A.; Marques, C. Gel-assisted formation of giant unilamellar vesicles. Biophysical journal 2013, 105 (1), 154–164.

(13) Starke-Peterkovic, T.; Turner, N.; Vitha, M. F.; Waller, M. P.; Hibbs, D. E.; Clarke, R. J. Cholesterol effect on the dipole potential of lipid membranes. Biophysical journal 2006, 90 (11), 4060–4070.

(14) Mondal, S.; Sarkar, M. Non-steroidal anti-inflammatory drug induced membrane fusion: concentration and temperature effects. The Journal of Physical Chemistry B 2009, 113 (51), 16323–16331. Struck, D. K.; Hoekstra, D.; Pagano, R. E. Use of resonance energy transfer to monitor membrane fusion. Biochemistry 1981, 20 (14), 4093–4099.

(15) O’donoghue, E. J.; Krachler, A. M. Mechanisms of outer membrane vesicle entry into host cells. Cellular microbiology 2016, 18 (11), 1508–1517.

(16) Greenspan, P.; Mayer, E. P.; Fowler, S. D. Nile red: a selective fluorescent stain for intracellular lipid droplets. The Journal of cell biology 1985, 100 (3), 965–973.

(17) Wang, X.; Eagen, W. J.; Lee, J. C. Orchestration of human macrophage NLRP3 inflammasome activation by Staphylococcus aureus extracellular vesicles. Proceedings of the National Academy of Sciences 2020.

(18) Van Meer, G.; Voelker, D. R.; Feigenson, G. W. Membrane lipids: where they are and how they behave. Nature reviews Molecular cell biology 2008, 9 (2), 112. Ingólfsson, H. I.; Carpenter, T. S.; Bhatia, H.; Bremer, P.-T.; Marrink, S. J.; Lightstone, F. C. Computational lipidomics of the neuronal plasma membrane. Biophysical journal 2017, 113 (10), 2271–2280.

(19) O’Donoghue, E. J.; Sirisaengtaksin, N.; Browning, D. F.; Bielska, E.; Hadis, M.; Fernandez-Trillo, F.; Alderwick, L.; Jabbari, S.; Krachler, A. M. Lipopolysaccharide structure impacts the entry kinetics of bacterial outer membrane vesicles into host cells. PLoS pathogens 2017, 13 (11), e1006760.

(20) Yang, S.-T.; Kiessling, V.; Tamm, L. K. Line tension at lipid phase boundaries as driving force for HIV fusion peptide-mediated fusion. Nature communications 2016, 7, 11401.

(21) Bandari, S.; Chakraborty, H.; Covey, D. F.; Chattopadhyay, A. Membrane dipole potential is sensitive to cholesterol stereospecificity: implications for receptor function. Chemistry and physics of lipids 2014, 184, 25–29.

(22) Gross, E.; Bedlack Jr, R. S.; Loew, L. M. Dual-wavelength ratiometric fluorescence measurement of the membrane dipole potential. Biophysical journal 1994, 67 (1), 208–216.

(23) Brockman, H. Lipid monolayers: why use half a membrane to characterize protein-membrane interactions? Current opinion in structural biology 1999, 9 (4), 438–443.

(24) Allende, D.; Vidal, A.; McIntosh, T. J. Jumping to rafts: gatekeeper role of bilayer elasticity. Trends in biochemical sciences 2004, 29 (6), 325–330. Brown, R. E.; Brockman, H. L. Using monomolecular films to characterize lipid lateral interactions. In Lipid Rafts, Springer, 2007; pp 41–58. Smaby, J. M.; Kulkarni, V. S.; Momsen, M.; Brown, R. E. The interfacial elastic packing interactions of galactosylceramides, sphingomyelins, and phosphatidylcholines. Biophysical journal 1996, 70 (2), 868–877.

(25) Feng, S.-s Interpretation of mechanochemical properties of lipid bilayer vesicles from the equation of state or pressure− area measurement of the monolayer at the air− water or oil− water interface. Langmuir 1999, 15 (4), 998–1010. Pinto, O. A.; Disalvo, E. A. A new model for lipid monolayer and bilayers based on thermodynamics of irreversible processes. Plos one 2019, 14 (4), e0212269. Auerswald, A.; Gruber, T.; Balbach, J.; Meister, A. Lipid-Dependent Interaction of Human N-BAR Domain Proteins with Sarcolemma Mono-and Bilayers. Langmuir 2020, 36 (30), 8695–8704.

(26) Goodrich, F. Proceedings, 2nd International Congress on Surface Activity. Vol. I 1957, 85.

(27) Peetla, C.; Jin, S.; Weimer, J.; Elegbede, A.; Labhasetwar, V. Biomechanics and thermodynamics of nanoparticle interactions with plasma and endosomal membrane lipids in cellular uptake and endosomal escape. Langmuir 2014, 30 (25), 7522–7532.

(28) Lewis, B. A.; Engelman, D. M. Lipid bilayer thickness varies linearly with acyl chain length in fluid phosphatidylcholine vesicles. Journal of molecular biology 1983, 166 (2), 211–217.

(29) Miyoshi, T.; Kato, S. Detailed analysis of the surface area and elasticity in the saturated 1, 2-diacylphosphatidylcholine/cholesterol binary monolayer system. Langmuir 2015, 31 (33), 9086–9096.

(30) Zhao, L.; Feng, S.-S. Effects of lipid chain length on molecular interactions between paclitaxel and phospholipid within model biomembranes. Journal of colloid and interface science 2004, 274 (1), 55–68.

(31) Simons, K.; Ikonen, E. Functional rafts in cell membranes. nature 1997, 387 (6633), 569.

(32) Yang, S.-T.; Kiessling, V.; Simmons, J. A.; White, J. M.; Tamm, L. K. HIV gp41– mediated membrane fusion occurs at edges of cholesterol-rich lipid domains. Nature chemical biology 2015, 11 (6), 424.

(33) Ermakov, Y. A.; Averbakh, A. Z.; Yusipovich, A. I.; Sukharev, S. Dipole potentials indicate restructuring of the membrane interface induced by gadolinium and beryllium ions. Biophysical journal 2001, 80 (4), 1851–1862.

(34) Chen, D.; Santore, M. M. Large effect of membrane tension on the fluid–solid phase transitions of two-component phosphatidylcholine vesicles. Proceedings of the National Academy of Sciences 2014, 111 (1), 179–184.

(35) Ge, J.; Shao, F. Manipulation of host vesicular trafficking and innate immune defence by Legionella Dot/Icm effectors. Cellular microbiology 2011, 13 (12), 1870–1880. Hubber, A.; Roy, C. R. Modulation of host cell function by Legionella pneumophila type IV effectors. Annual review of cell and developmental biology 2010, 26, 261–283.

(36) Asawakarn, T.; Cladera, J.; O’Shea, P. Effects of the membrane dipole potential on the interaction of saquinavir with phospholipid membranes and plasma membrane receptors of Caco-2 cells. Journal of Biological Chemistry 2001, 276 (42), 38457–38463. Samsonov, A. V.; Chatterjee, P. K.; Razinkov, V. I.; Eng, C. H.; Kielian, M.; Cohen, F. S. Effects of membrane potential and sphingolipid structures on fusion of Semliki Forest virus. Journal of virology 2002, 76 (24), 12691–12702. Wang, L.; Bose, P. S.; Sigworth, F. J. Using cryo-EM to measure the dipole potential of a lipid membrane. Proceedings of the National Academy of Sciences 2006, 103 (49), 18528–18533.

(37) Nomura, K.; Inaba, T.; Morigaki, K.; Brandenburg, K.; Seydel, U.; Kusumoto, S. Interaction of lipopolysaccharide and phospholipid in mixed membranes: solid-state 31P-NMR spectroscopic and microscopic investigations. Biophysical journal 2008, 95 (3), 1226–1238.

(38) Augenstreich, J.; Haanappel, E.; Ferré, G.; Czaplicki, G.; Jolibois, F.; Destainville, N.; Guilhot, C.; Milon, A.; Astarie-Dequeker, C.; Chavent, M. The conical shape of DIM lipids promotes Mycobacterium tuberculosis infection of macrophages. Proceedings of the National Academy of Sciences 2019, 116 (51), 25649–25658.

(39) Mura, M.; Dennison, S. R.; Zvelindovsky, A. V.; Phoenix, D. A. Aurein 2.3 functionality is supported by oblique orientated α-helical formation. Biochimica et Biophysica Acta (BBA)-Biomembranes 2013, 1828 (2), 586–594.

(40) Adams, P. G.; Lamoureux, L.; Swingle, K. L.; Mukundan, H.; Montaño, G. A. Lipopolysaccharide-induced dynamic lipid membrane reorganization: tubules, perforations, and stacks. Biophysical journal 2014, 106 (11), 2395–2407.

(41) Szule, J. A.; Fuller, N. L.; Rand, R. P. The effects of acyl chain length and saturation of diacylglycerols and phosphatidylcholines on membrane monolayer curvature. Biophysical journal 2002, 83 (2), 977–984.

(42) Risselada, H. J.; Marelli, G.; Fuhrmans, M.; Smirnova, Y. G.; Grubmüller, H.; Marrink, S. J.; Müller, M. Line-tension controlled mechanism for influenza fusion. PLoS One 2012, 7 (6), e38302.

(43) Zhou, Y.; Wong, C.-O.; Cho, K.-j.; Van Der Hoeven, D.; Liang, H.; Thakur, D. P.; Luo, J.; Babic, M.; Zinsmaier, K. E.; Zhu, M. X. Membrane potential modulates plasma membrane phospholipid dynamics and K-Ras signaling. Science 2015, 349 (6250), 873–876.

(44) Kovács, T.; Batta, G.; Hajdu, T.; Szabó, Á.; Váradi, T.; Zákány, F.; Csomós, I.; Szöllősi, J.; Nagy, P. The dipole potential modifies the clustering and ligand binding affinity of ErbB proteins and their signaling efficiency. Scientific reports 2016, 6, 35850.

(45) Nitenberg, M.; Benarouche, A.; Maniti, O.; Marion, E.; Marsollier, L.; Géan, J.; Dufourc, E. J.; Cavalier, J.-F.; Canaan, S.; Girard-Egrot, A. P. The potent effect of mycolactone on lipid membranes. PLoS pathogens 2018, 14 (1), e1006814.

(46) Lee, E. Y.; Bang, J. Y.; Park, G. W.; Choi, D. S.; Kang, J. S.; Kim, H. J.; Park, K. S.; Lee, J. O.; Kim, Y. K.; Kwon, K. H. Global proteomic profiling of native outer membrane vesicles derived from Escherichia coli. Proteomics 2007, 7 (17), 3143–3153. Scorza, F. B.; Doro, F.; Rodríguez-Ortega, M. J.; Stella, M.; Liberatori, S.; Taddei, A. R.; Serino, L.; Moriel, D. G.; Nesta, B.; Fontana, M. R. Proteomic characterization of outer membrane vesicles from the extraintestinal pathogenic Escherichia coli tolR IHE3034 mutant. Molecular & Cellular Proteomics 2007.

