## Supporting Information for "Lipid Specificity of the Fusion of Bacterial Extracellular Vesicles with the Host Membrane"

[#] Current Affiliation

† Equal authorship

* Corresponding author: Mohammed Saleem.

Contact: +91-674-2494501

Running Title: Bacterial MV – Host Membrane Interaction

**Theoretical Estimation of bacterial MVs Concentration:**

Purified MV concentration as well as lipid estimation of isolated bacterial MVs were calculated as follows:

Prior to calculation it was assumed that (i) the dispersion had only unilamellar vesicles and (ii) all MVs were of the equal size as evident from the DLS and electron micrographs (Fig 1B and 1D).

Number of phospholipid molecules per MV, $N_{molecules} = \frac{[4\pi R^2 + 4\pi\,(R-h)^2]}{a}$

Where,

$R$ = Average radius of each vesicle= 100 nm (From Fig 1C),

$h$ = Average thickness of bilayer (5 nm for DSPC) and

$a$ = Cross sectional area of a lipid head-group supposed to be 0.6 $nm^2$ (estimated for PC headgroup).

Considering the above values, $N_{molecules}$= 398256.

Number of MVs per ml, $N_{MV} = \frac{M \times N_A}{N_{molecules} \times 1000}$

where, $M$= Molar mass of each lipid (800 g/mol), $N_A$ is Avogadro's number.

Since,

$N_{MV} = 1.21 \times 10^{18}$.

Therefore,

Moles of MV= $\frac{N_{MV}}{N_A}$

Hence,

2µM MV in the suspension.

**Fig. S1**

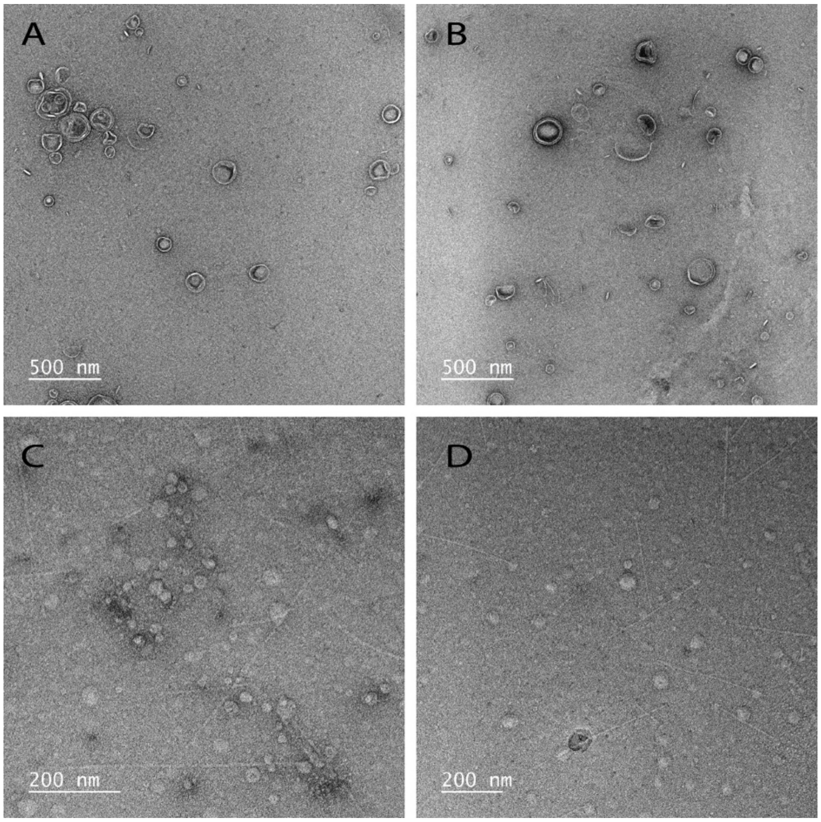


**E**

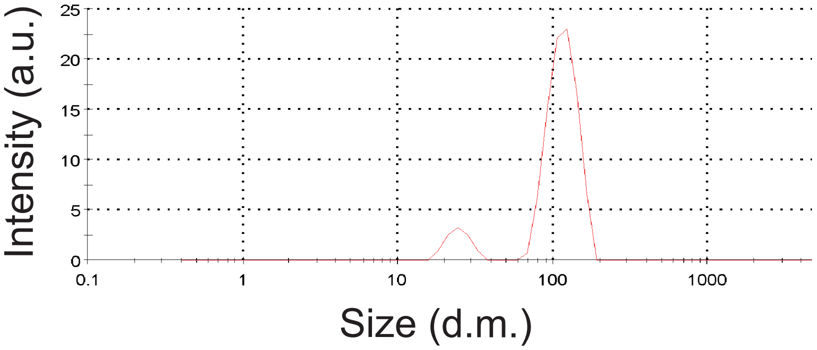


**Fig. S1: Electron micrograph of MVs isolated by ultracentrifugation (A and B) and chemical vesiculation method (C and D). E is the size distribution of MVs isolated by chemical vesiculation method.**

Fig. S2

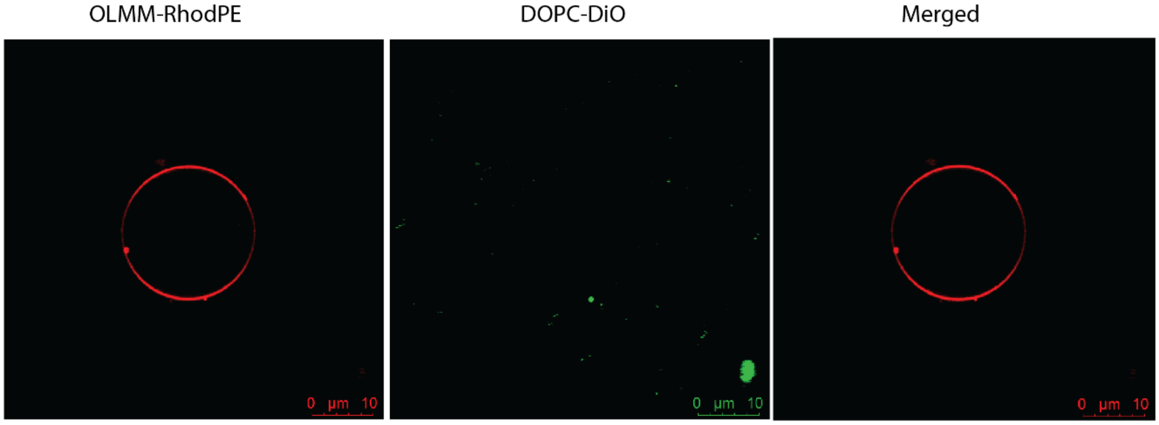


**Fig. S2. Di-O labelled LUVs do not fuse with outer leaflet model membrane (OLMM).**

**Fig. S3**

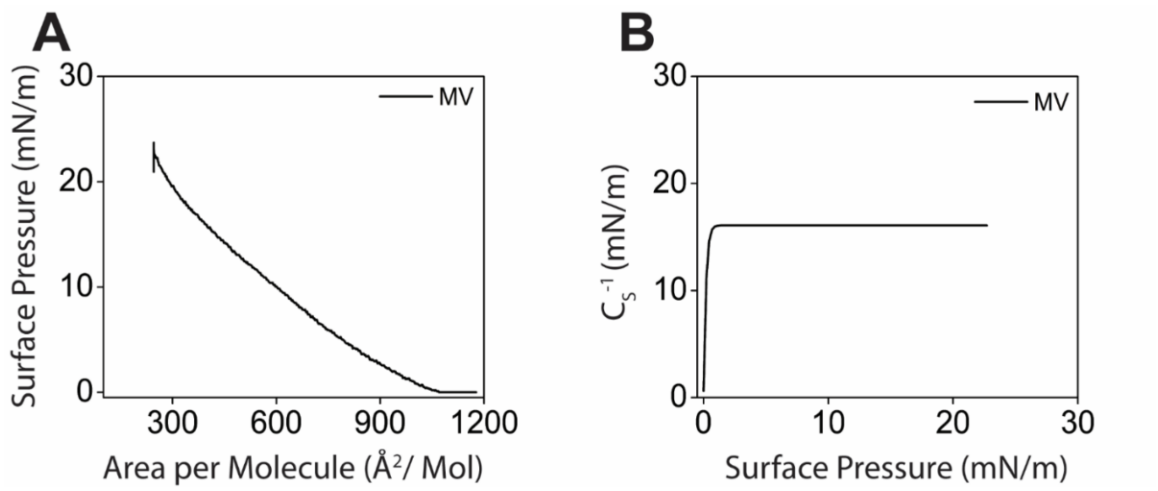


**Fig. S3. (A) Surface Pressure (π) - Mean molecular area (A) isotherm of bacterial MV and (B) Compressibility modulus (in-plane elasticity, $C_s^{-1}$) with respect to surface pressure of MV.**

**Fig. S4**

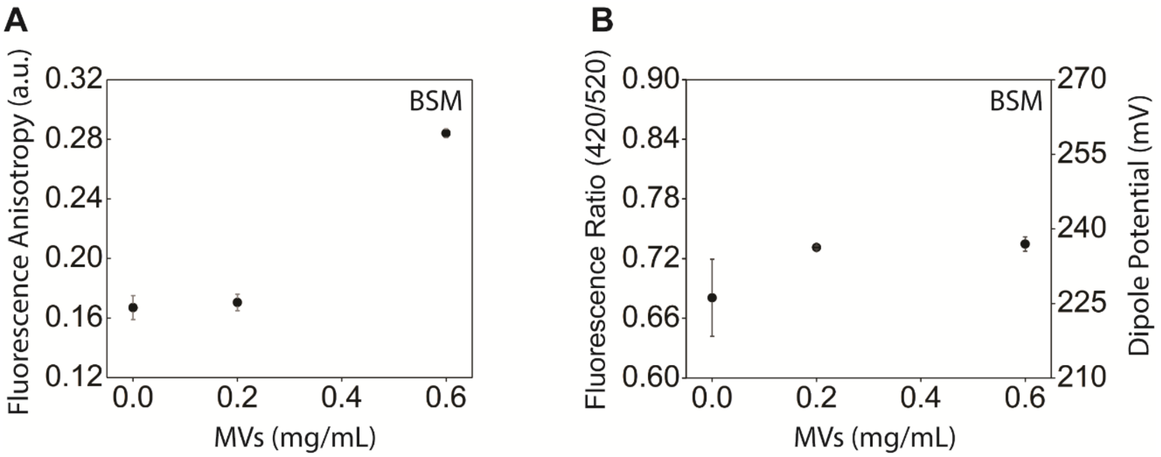


**Fig. S4. Change in (A) anisotropy and (B) fluorescence ratio/dipole potential of BSM membrane incubates with bacterial MVs. Data points are shown as mean ± SE, n = 3.**

**Fig. S5**

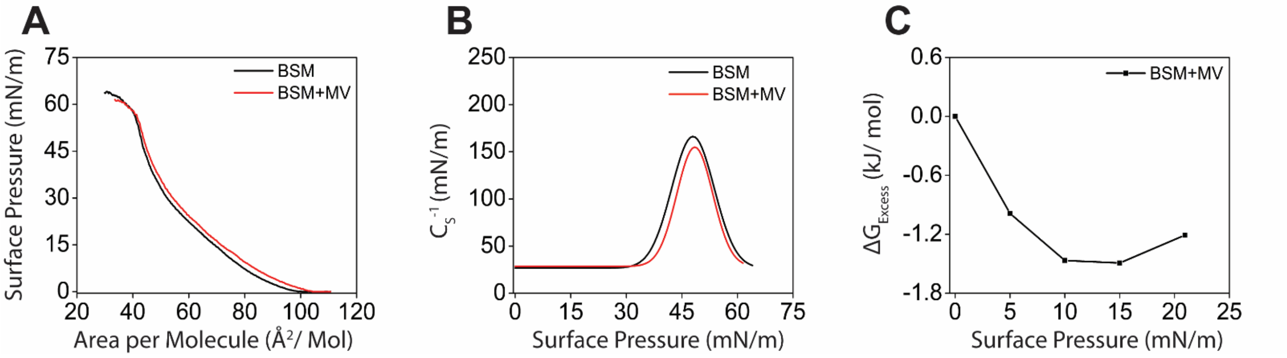


**Fig. S5. (A) Plot showing Surface Pressure (π) - Mean molecular area (A) isotherm of BSM (black curve) as control and BSM incubated with bacterial MVs (red curve) (B) Compressibility modulus (in-plane elasticity, $C_s^{-1}$) with respect to surface pressure of BSM (black curve) and BSM with MV (red curve) (C) Excess Gibb's free energy of mixing between BSM and bacterial MVs.**

**Fig. S6**

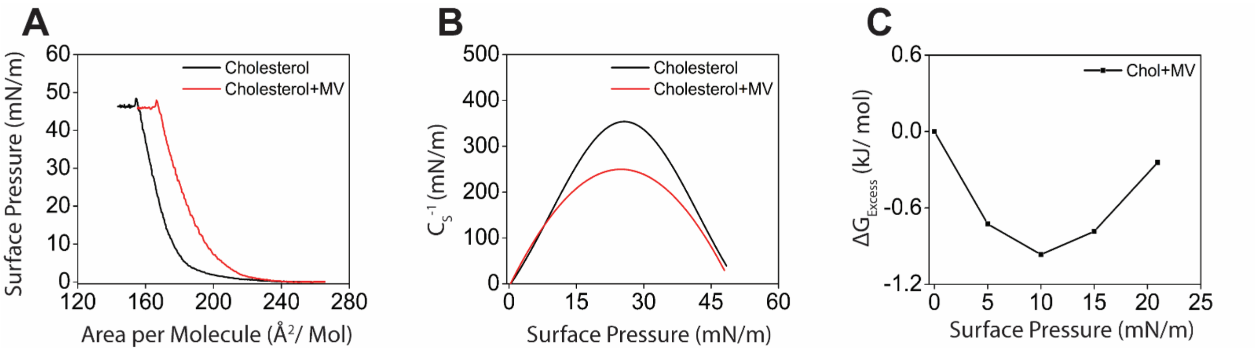


**Fig. S6. (A) Plot showing Surface Pressure (π) - Mean molecular area (A) isotherm of cholesterol (black curve) as control and cholesterol incubated with bacterial MVs (red curve) (B) Compressibility modulus (in-plane elasticity, $C_s^{-1}$) with respect to surface pressure of cholesterol (black curve) and cholesterol with MV (red curve) (C) Excess Gibb's free energy of mixing between cholesterol and bacterial MVs.**

**Fig. S7.**

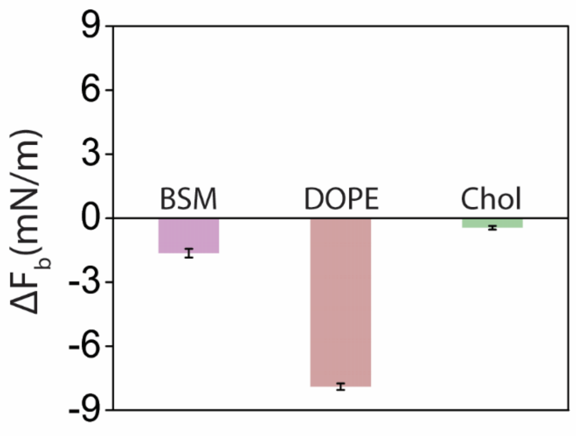


**Fig. S7. Fig. showing histogram depicting change in bending force ($\Delta F_b$) of BSM, DOPE and Cholesterol monolayers at the interface in presence of bacterial MVs. Data points shown as mean ± SE, n = 3.**

**Fig S8.**

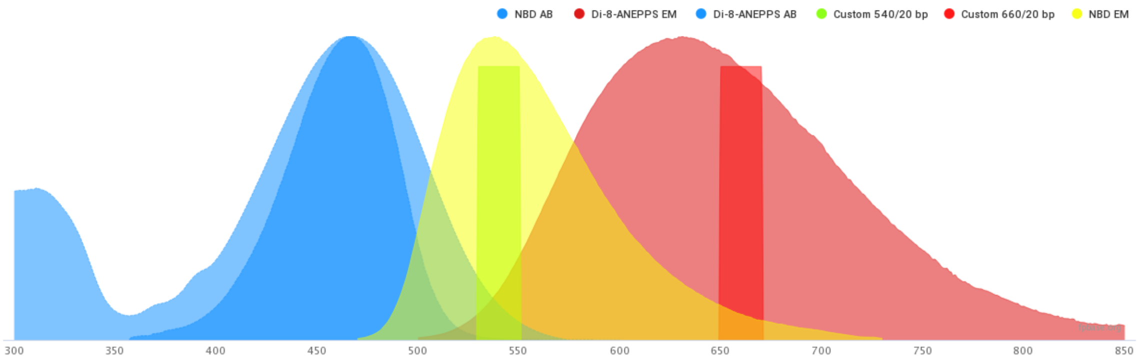


Collection Efficiency (%)

| Filter | Di-8-ANEPPS | NBD |
|---|---|---|
| Custom 540/20 bp | 1.9 | 19.4 |
| Custom 660/20 bp | 10.5 | 1.7 |

**Fig S8. Showing the set-up for confocal microscopy for NBD-PE and Di-8-ANEPPS dyes.**

**Table S1. Surface potential of phospholipid membranes and bacterial membrane vesicles (MVs)**

| **Phospholipid/MV** | **Zeta Potential (mV)** |
|---|---|
| Bacterial Membrane Vesicles (MVs) | -9.5 |
| DOPC | -18.6 |

**Table S2. Table showing change in bending force induced by bacterial MVs in monolayers of different monolayer compositions. Data recorded as mean ± SE, n = 3.**

| **Monolayer composition** | **$\Delta F_b$ (mN/m)** |
|---|---|
| OLMM | -1.75 ± 0.13 |
| DOPC | -1.52 ± 0.12 |
| DLPC | -3.15 ± 0.03 |
| DMPC | -5.04 ± 0.23 |
| DPPC | -7.16 ± 0.1 |
| DSPC | -1.5 ± 0.26 |
| 5:5:0 | -1.37 ± 0.01 |
| 4:4:2 | 0.44 ± 0.06 |
| 3:3:4 | -0.7 ± 0.17 |
| BSM | -1.64 ± 0.2 |
| DOPE | -7.9 ± 0.15 |
| Cholesterol | -0.45 ± 0.08 |

**Table S3. Table showing excess Gibb's free energy of mixing ($\Delta G_{Excess}$) and mixing enthalpy ($\Delta H$) of bacterial MVs with lipid monolayers of different compositions at various surface pressures.**

| Surface Pressure $\pi$, (mN/m) | $\pi$ = 5 mN/m | | | $\pi$ = 10 mN/m | | | $\pi$ = 15mN/m | | | $\pi$ = 20.95 mN/m | | |
|---|---|---|---|---|---|---|---|---|---|---|---|---|
| Monolayer composition | $\Delta G_{Excess}$ (kJ/mol) | $\Delta G_{Excess}$ (kT) | $\Delta H$ (kJ/mol) | $\Delta G_{Excess}$ (kJ/mol) | $\Delta G_{Excess}$ (kT) | $\Delta H$ (kJ/mol) | $\Delta G_{Excess}$ (kJ/mol) | $\Delta G_{Excess}$ (kT) | $\Delta H$ (kJ/mol) | $\Delta G_{Excess}$ (kJ/mol) | $\Delta G_{Excess}$ (kT) | $\Delta H$ (kJ/mol) |
| OLMM | -0.733 | -0.295 | -2.094 | -1.061 | -0.428 | -3.03 | -1.014 | -0.409 | -2.895 | -0.709 | -0.286 | -2.025 |
| DOPC | -0.764 | -0.308 | -2.347 | -1.161 | -0.468 | -3.568 | -1216 | -0.49 | -3.739 | -1038 | -0.418 | -3.191 |
| DLPC | -0.417 | -0.168 | -1.867 | -0.598 | -0.241 | -2.68 | -0.572 | -0.23 | -2.565 | -0.435 | -0.175 | -1.949 |
| DMPC | -0.437 | -0.176 | -2.03 | -0.634 | -0.255 | -2.943 | -0.627 | -0.252 | -2.906 | -0.473 | -0.191 | -1.096 |
| DPPC | -0.903 | -0.364 | 2.317 | -1.356 | -0.546 | -3.479 | -1.396 | -0.563 | -3.582 | -1.169 | -0.471 | -1.5 |
| DSPC | -0.57 | -0.229 | -2.093 | -0.832 | -0.335 | -3.067 | -0.821 | -0.331 | -1.506 | -0.628 | -0.253 | -1.153 |
| 5:5:0 | -0.937 | -0.377 | -2.314 | -1.345 | -0.542 | -3.32 | -1.291 | -0.52 | -1.594 | -0.921 | -0.371 | -1.136 |
| 4:4:2 | -0.539 | -0.217 | -1.462 | -0.688 | -0.277 | -1.811 | -0.418 | -0.168 | -1.134 | 0.129 | 0.052 | 0.174 |
| 3:3:4 | -1.205 | -0.486 | -2.628 | -1.785 | -0.72 | -3.893 | -1.757 | -0.708 | -3.834 | -1432 | -0.577 | -1.561 |
| SM | -0.988 | -0.398 | -2.424 | -1.466 | -0.591 | -3.6 | -1.492 | -0.601 | -3.663 | 1.208 | -0.487 | -2.966 |
| DOPE | -0.686 | -0.276 | -2.181 | -0.987 | -0.389 | -3.138 | -0.952 | -0.316 | -1.513 | -0.697 | -0.098 | -1.107 |
| Cholesterol | -0.726 | -0.292 | -0.911 | -0.966 | -0.398 | -1.214 | -0.784 | -0.384 | -0.985 | -0.243 | -0.281 | -0.306 |

### SI References


1. Sezgin E*, et al.* (2012) Elucidating membrane structure and protein behavior using giant plasma membrane vesicles. *nature protocols* 7(6):1042.
2. Folch J, Lees M, & Sloane Stanley G (1957) A simple method for the isolation and purification of total lipides from animal tissues. *J biol Chem* 226(1):497-509.
3. Reis A*, et al.* (2013) A comparison of five lipid extraction solvent systems for lipidomic studies of human LDL. *Journal of lipid research* 54(7):1812-1824.
4. Starke-Peterkovic T*, et al.* (2006) Cholesterol effect on the dipole potential of lipid membranes. *Biophysical journal* 90(11):4060-4070.
